# Hypothalamic NMDA receptors stabilize NREM sleep and are essential for REM sleep

**DOI:** 10.1101/2020.10.19.345728

**Authors:** Giulia Miracca, Berta Anuncibay Soto, Kyoko Tossell, Raquel Yustos, Alexei L. Vyssotski, Nicholas P. Franks, William Wisden

## Abstract

The preoptic hypothalamus regulates both NREM and REM sleep. We found that calcium levels in mouse lateral preoptic (LPO) neurons were highest during REM. Deleting the core GluN1 subunit of NMDA receptors from LPO neurons abolished calcium signals during all vigilance states, and the excitatory drive onto LPO neurons was reduced. Mice had less NREM sleep and were incapable of generating conventionally classified REM sleep episodes: cortical theta oscillations were greatly reduced but muscle atonia was maintained. Additionally, mice lacking NMDA receptors in LPO neurons had highly fragmented sleep-wake patterns. The fragmentation persisted even under high sleep pressure produced by sleep deprivation. Nevertheless, the sleep homeostasis process remained intact, with an increase in EEG delta power. The sedative dexmedetomidine and sleeping medication zolpidem could transiently restore consolidated sleep. High sleep-wake fragmentation, but not sleep loss, was also produced by selective GluN1 knock-down in GABAergic LPO neurons. We suggest that NMDA glutamate receptor signalling stabilizes the firing of “GABAergic NREM sleep-on” neurons and is also essential for the theta rhythm in REM sleep.

## INTRODUCTION

Many people suffer from occasional insomnia, but it can develop into a debilitating condition (Roth et al., 2011). Insomnia, as a clinical disorder, is defined as an inability to initiate or maintain sleep at least three times a week over three months, even when sleep conditions are otherwise optimal (Van Someren, 2020). Insomniacs frequently report that their sleep is non-restorative and that they sleep less. In fact, insomnia sufferers often have the same amounts of EEG-defined NREM sleep as controls, but oscillate frequently between wake and NREM sleep, so that their sleep is fragmented, and often their REM sleep is disturbed (Van Someren, 2020). For self-reported sleep quality, good sleep continuity and increased length REM sleep were found to give the best subjective sleep quality (Della Monica et al., 2018). Even if sleep is poor, wakefulness still cannot be sustained beyond a certain limit. This limit is thought to be imposed by the process of sleep homeostasis, also named Process S, the increasing drive to enter NREM sleep as wakefulness continues (Borbely et al., 2016). S then declines as sleep progresses.

Both NREM and REM sleep are in part controlled by the preoptic (PO) hypothalamus (Lu et al., 2002; Lu et al., 2000; McGinty and Sterman, 1968; Nauta, 1946; Sherin et al., 1996; Szymusiak et al., 2007). In this structure, GABA/peptidergic neurons, e.g. GABA/galanin neurons, contribute to NREM sleep induction and sleep homeostasis (Chung et al., 2017; Kroeger et al., 2018; Ma et al., 2019; Reichert et al., 2019; Sherin et al., 1996; Zhang et al., 2015). To stay asleep, and prevent insomnia, it seems reasonable to assume that these sleep-promoting neurons would have to stay “on”. Indeed, lesioning of lateral (LPO) neurons in rats reduces the amounts of NREM or REM sleep, depending on the lesion’s location (Lu et al., 2000). But factors that keep LPO sleep-promoting neurons firing and so govern the lengths of NREM and REM sleep episodes are not known.

In both insects and mammals, calcium entry through NMDA-type ionotropic glutamate-gated receptors has been proposed to signal the sleep homeostatic process, in an analogous mechanism to changing synaptic strength that underlies memory formation (Liu et al., 2016; Raccuglia et al., 2019; Tatsuki et al., 2016). These studies provided our initial motivation for the experiments reported here. We wanted to test if NMDA receptors mediate the homeostatic sleep drive in the PO hypothalamus. Consistent with the hypothesis, NMDA receptor activation promotes sleep. In the fruit fly *Drosophila*, genetic knockdown of NMDA receptors in brain reduces total sleep time (Tomita et al., 2015). In rodents, NMDA receptor antagonists reduce *and* agonists enhance NREM sleep (Burgdorf et al., 2019; Tatsuki et al., 2016). Furthermore, patients with autoimmunity to the core and essential GluN1 subunit of NMDA receptors often suffer severe insomnia (Arino et al., 2020; Dalmau et al., 2019). Because NMDA receptors are near universally expressed in the brain (Monyer et al., 1994; Moriyoshi et al., 1991), these effects on sleep could come from interference with many circuits.

In this study, we deleted the GluN1 NMDA receptor subunit in the LPO hypothalamus and find that NMDA receptors there are not involved in sleep homeostasis. Instead, we obtained an unexpected “insomnia” phenotype with high sleep-wake fragmentation and greatly diminished REM sleep. This implicates a new signaling pathway in maintaining the consistency of NREM sleep and generating REM sleep. The sleep-wake fragmentation effect is selective for the GluN1 expression in GABA neurons. We suggest that NMDA glutamate receptor signalling, perhaps because of the long open times of this type of channel, stabilizes the firing of hypothalamic “GABAergic sleep-on” neurons for consolidated sleep.

## RESULTS

### CALCIUM ACTIVITY IN LPO HYPOTHALAMIC NEURONS IS HIGHEST DURING REM SLEEP

We first recorded calcium activity in LPO hypothalamic neurons using photometry with GCaMP6s expressed under the control of a universally active promoter. Mice were injected in LPO with *AAV-GCaMP6s* (Figure 1A-B). Highest calcium activity occurred during REM sleep episodes, especially at the beginning and end of the episodes (Figure 1C-D). During NREM sleep, LPO neurons showed a more sporadic and spikey activity, and during wakefulness only low activity. By plotting scaled means of GCaMP6s signal against EMG signal (Figure 1E, left panel) and delta power (Figure 1E, centre panel), REM sleep episodes were distributed towards higher values of GCaMP6s signal, separating them from the NREM sleep and wake data points. Comparing the average of ΔF/F points during each behavioural state, REM sleep calcium values were significantly higher than in other vigilance states (Figure 1E, right panel).

**Figure 1.**
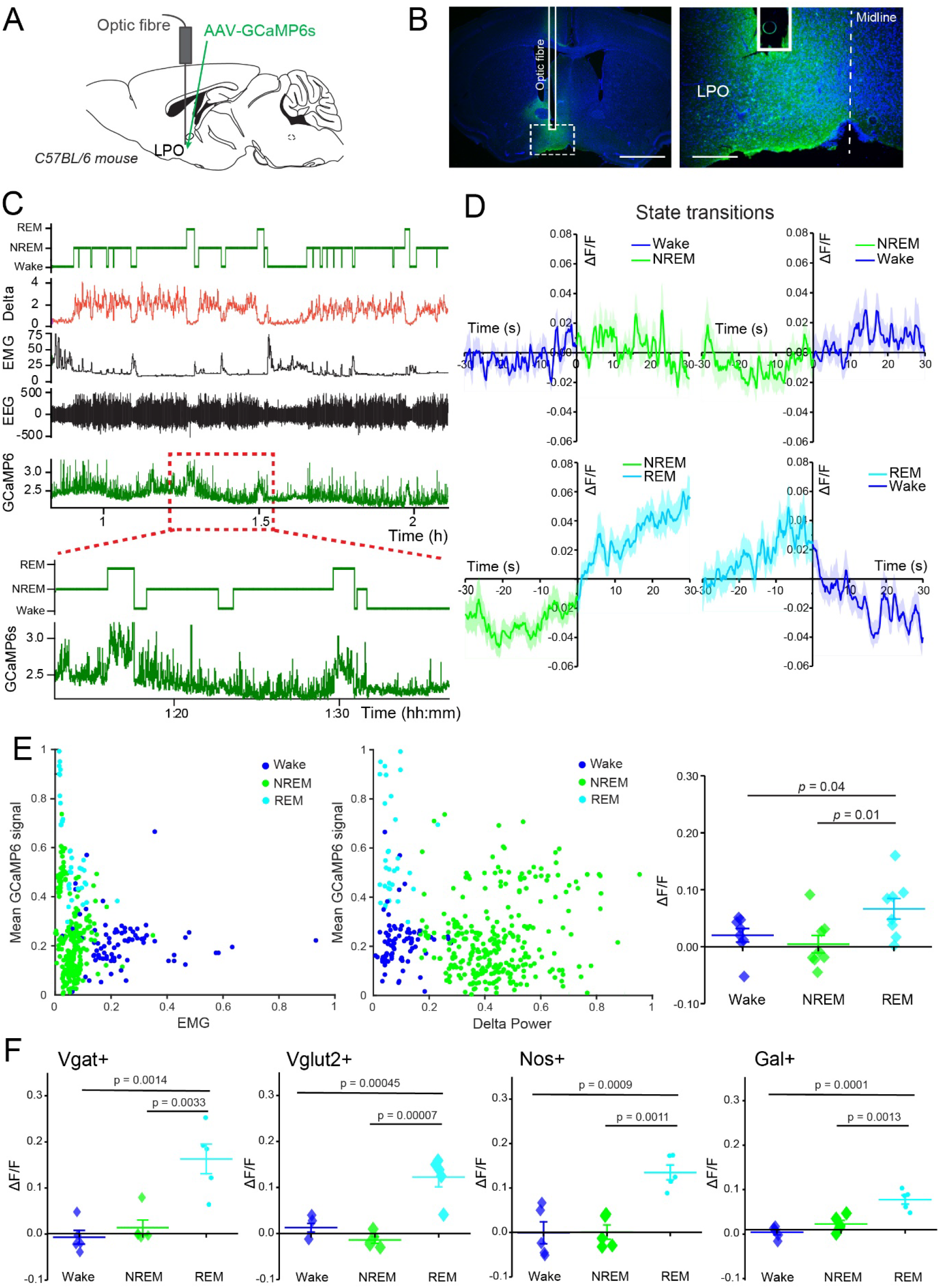
Calcium activity in LPO hypothalamic neurons is highest during REM sleep. **A**, schematic representation of optic fibre implantation and *AAV-GCaMP6s* injection in LPO area of C57BL/6 mice. **B**, immunohistochemistry staining to show optic fibre track and viral vector expression in LPO, using green fluorescent protein (GFP, in green) antisera and 4’,6-diamidino-2-phenylindole (DAPI, in blue). Scale bars left 1 mm, right 200 μm. **C**, example of photometry recording from LPO neurons aligned to EEG and EMG data. From the top: stage, delta power, EMG, EEG and GCaMP6s signal. The dotted red square indicates the trace’s segment expanded below. **D**, photometry recordings across state transitions normalized as ΔF/F data points. For the transitions to be considered, animals had to be in the behavioural state before and after transitions for at least 30s each. **E**, left and centre panels, scaled mean of GCaMP6s signal plotted against EMG (left) and delta power (right); right panel, quantification of LPO calcium activity as ΔF/F by behavioural state (n = 8 mice, F = 4.473, *P*= 0241). **F**, screening of LPO neuronal populations for calcium activity shown as ΔF/F using gene-specific Cre mouse lines and injection of *AAV-flex-GCaMP6s*. From the left: Vgat-Cre, (*n* = 5 mice, F = 16.87, *P* = 0.0003), Vglut2-Cre (*n* = 5 mice, F = 26.94, *P* < 0.0001), Nos1-Cre (*n* = 5 mice, F = 15.82, *P* = 0.0004) and Galanin-Cre (*n* = 5 mice, F = 21.55, *P* = 0.0001). In D, E and F data are represented as means ± SEM. In E and F, significance was calculated using one-way ANOVA followed by Tukey’s *post-hoc* test was used. P values showed in these plots are obtained from multiple comparisons analysis and corrected by Tukey’s *post-hoc* test.

Photometry recordings were also performed in different subtypes of LPO hypothalamic neurons by injecting *AAV-flex-GCaMP6s* into the LPO area of *Vgat-Cre*, *Vglut2-Cre*, *Nos1-Cre* and *Galanin-Cre* mice (Supplementary Figure 1A-C). As for the pan-neuronal recordings, the subsets of LPO neuronal populations all showed a significantly higher calcium activity during REM sleep episodes compared with NREM sleep and wake (Figure 1F).

### DELETION OF NMDA RECEPTORS FROM LPO NEURONS ABOLISHES CALCIUM SIGNALS

NMDA receptors assemble as heteromeric tetramers of subunits, with two core GluN1 subunits, whose gene *grin1* is transcribed universally in the brain (Moriyoshi et al., 1991), and GluN2 and/or GluN3 subunits, whose genes are differentially expressed (Monyer et al., 1994; Monyer et al., 1992; Paoletti et al., 2013). GluN1 is essential for all NMDA receptors (Paoletti et al., 2013; Tsien et al., 1996). Mice homozygous for a conditional allele (floxed-*Grin1*) that encodes the GluN1 subunit were bilaterally injected into the LPO hypothalamus with *AAV-Cre-2A-Venus*, generating ΔGluN1-LPO mice (Figure 2A). More than 75% of the cells expressing the *AAV-Cre-2A-Venus* transgene were in LPO (Supplementary Figure 2A-C). Microglia and astrocytes were not transduced with the particular AAV (AAV1/2) serotype, as there was no co-expression of Venus and the astrocyte marker GFAP or the microglial marker IBA1 (Supplementary Figure 2D).

**Figure 2.**
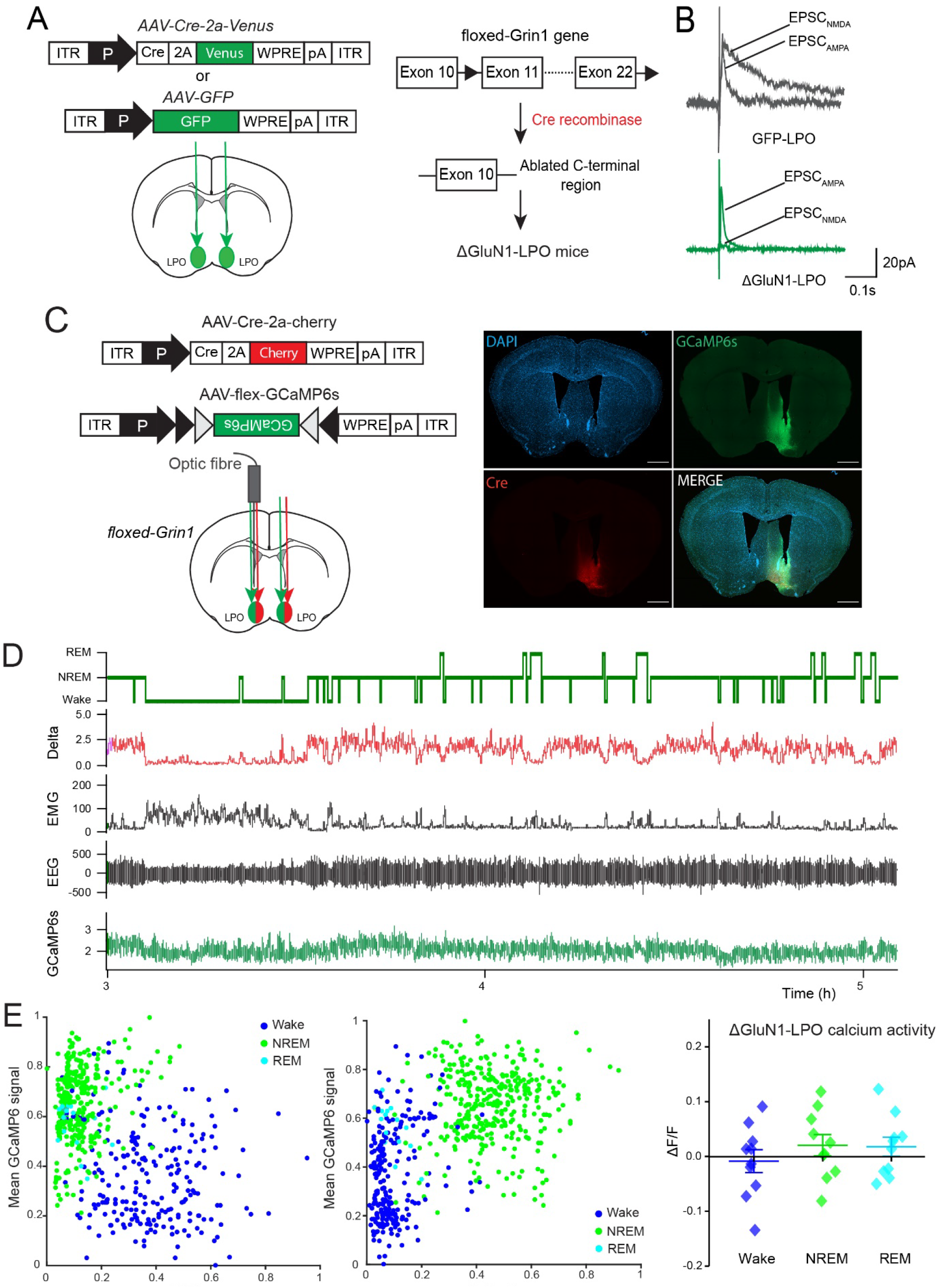
Calcium activity in LPO hypothalamic neurons requires NMDA receptors. **A**, generation of ΔGluN1-LPO and GFP-LPO animals by bilateral injection in LPO of *AAV-Cre-2A-Venus* and *AAV-GFP* respectively. **B**, evoked spontaneous post-synaptic currents (EPSCs) in LPO neurons to show NMDA and AMPA receptors currents in GFP-LPO animals (top) and the deletion of NMDA receptor currents in ΔGluN1-LPO cells (bottom). **C**, floxed-*Grin1* animals were bilaterally co-injected in LPO with *AAV-Cre-2a-cherry* and *AAV-flex-GCaMP6s* viral vectors to record calcium activity only from neurons lacking the NMDA receptors. An optic fibre was also implanted unilaterally for these recordings. On the right, immunohistochemistry showing from left to right: DAPI (blue) and *AAV-flex GCaMP6s* transgene (green) on the first row, Cre recombinase (red) and merge on the second row. Scale bars represent 1 mm. **D**, example of a photometry recording aligned to EEG and EMG in a ΔGluN1-LPO mouse. From the top: vigilance state, delta power, EMG, EEG and GCaMP6s signal. **E** left and centre panels, scaled mean of GCaMP6s signal plotted against EMG (left) and delta power (centre); right panel, quantification of ΔGluN1-LPO calcium activity normalized as ΔF/F during behavioural states. One-way ANOVA followed by Tukey’s *post-hoc* test (F = 0.6917, *P* = 5. 094). ΔGluN1-LPO, *n* = 5. Data in E (right panel) are represented as mean ± SEM.

We examined if NMDA receptor currents were deleted from ΔGluN1-LPO neurons compared to control mice injected with AVV-GFP by recording evoked excitatory post-synaptic currents (eEPSCs) from *ex-vivo* acute slices prepared from the PO area (Supplementary Figure 2E and F). eEPSCs on LPO neurons consist of both a slow (hundreds of milliseconds) NMDA and fast (few millisecond) AMPA receptor component (Figure 2B). On LPO neurons from ΔGluN1-LPO mice, the slow NMDA component was eliminated (Figure 2B). Deletion of the NMDA-mediated current from LPO neurons also reduced spontaneous excitatory post-synaptic current (sEPSCs) amplitudes (Supplementary Figure 2H) and frequency (Supplementary Figure 2I).

We tested how NMDA deletion from LPO neurons influenced intracellular calcium. We co-injected into the LPO area of *floxed-Grin1* gene mice *AAV-Cre-Cherry* and *AAV-flex - GCaMP6s*, so that only neurons expressing Cre recombinase express the calcium sensor (Figure 2C). Regardless of vigilance state, calcium activity in ΔGluN1-LPO neurons was greatly reduced (Figure 2D and E), and REM sleep was no longer highlighted by calcium activity, showing that deleting NMDA receptors ablates calcium fluctuations from LPO neurons.

### DELETION OF NMDA RECEPTORS FROM LPO NEURONS REDUCES NREM AND REM SLEEP AND PRODUCES HIGH SLEEP-WAKE FRAGMENTATION

We examined the effect of the GluN1 LPO deletion on vigilance states (Figure 3A). Throughout 24h, ΔGluN1-LPO mice compared with GFP-LPO (Figure 3B) mice spent more time awake (Figure 3C and D). ΔGluN1-LPO mice lost on average 15-20% of NREM sleep time and 50% of their REM sleep time during both light and dark periods (Figure 3D). EEG power spectra for wake and NREM were similar between control and ΔGluN1-LPO mice (Figure 3E); however, ΔGluN1-LPO mice had strongly reduced cortical theta oscillations during REM sleep episodes (5-10 Hz, Figure 3E, right panel). Consequently, EEG and EMG recordings during REM sleep episodes (Supplementary Figure 3A and B) showed a reduced theta:delta power (T:D) ratio in ΔGluN1-LPO mice compared to GFP-LPO controls (Supplementary Figure 3C), although atonia was maintained in both groups.

**Figure 3.**
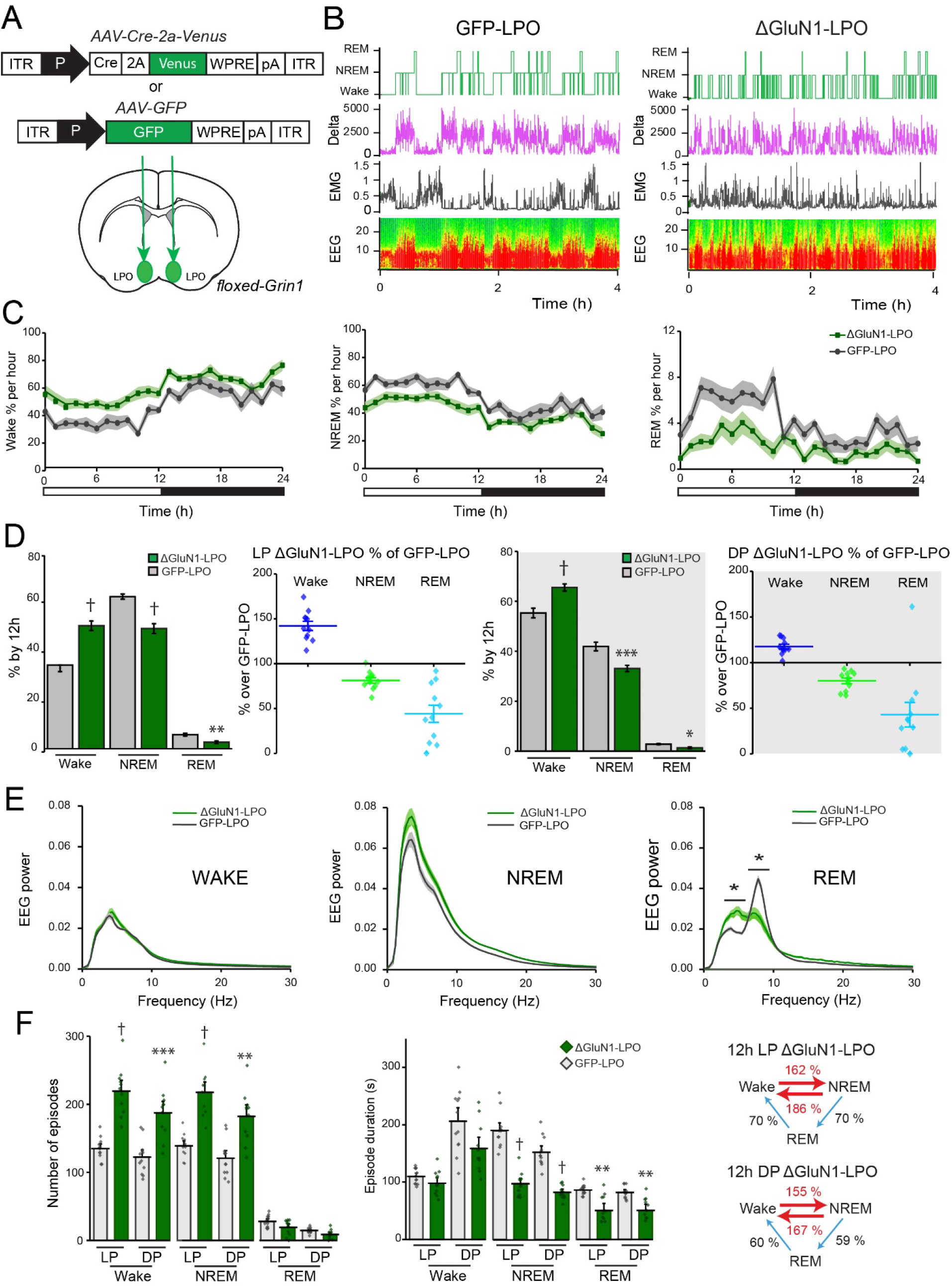
Deletion of NMDA receptors in LPO hypothalamus reduces NREM and REM sleep time and produces sleep-wake fragmentation. **A**, bilateral *AAV-Cre-Venus* and *AAV-GFP* virus injections into LPO of floxed-*Grin1* mice to generate ΔGluN1-LPO and GFP-LPO control animals respectively. **B**, example of baseline recordings from GFP-LPO (left) and ΔGluN1-LPO (right) animals. From the top, hypnogram, delta power, EMG, and EEG represented as a somnogram and as a trace. **C**, 24-hour baseline states distribution as percentage of 1h of wake (left), NREM (centre) and REM sleep (right) during light (0-12h) and dark (12-24h) period. 2-way repeated measures (RM) ANOVA followed by Sidak’s *post-hoc* test, using “time” and “virus” as factors. Wake, “time”: F (8.612, 180.9) = 14.01, P<0.0001. NREM, “time”: F (8.972, 188.4) = 12.67, P<0.0001; “virus”: F (1, 21) = 41.26, P<0.0001. REM, “time”: F (9.057, 190.2) = 8.307, P<0.0001; “virus”: F (1, 21) = 18.06, P=0.0004. **D** 1^st^ and 3^rd^ panels, quantification of each behavioural state as percentage over 12 hours of light (1^st^ panel) and dark period (3^rd^ panel). 2-way ANOVA followed by Sidak’s *post-hoc* test using “state” and “virus” as factors. LP, “state”: F (2, 63) = 795.5, P<0.0001; “virus”: F (1, 63) = 0.0002798, P=0.9867. DP, “state”: F (2, 63) = 956.6, P<0.0001; “virus”: F (1, 63) = 1.230e-005, P=0.9972; 2^nd^ and 4^th^ panels, wake, NREM and REM sleep amounts in ΔGluN1-LPO mice represented as percentage of GFP-LPO mice amounts during the light (2^nd^ panel) and dark (4^th^ panel) periods. **E**, EEG power spectrum for wake (left), NREM (centre) and REM sleep (right) normalized over total EEG power. 2-way ANOVA followed by Sidak’s *post-hoc* test using “frequency” and “virus” as factors. Wake “frequency”: F (89, 1800) = 745.9, P<0.0001; “virus”: F (1, 1800) = 9.142e-009, P>0.9999. NREM “frequency”: F (89, 1800) = 1601, P<0.0001; “virus”: F (1, 1800) = 1.424e-008, P>0.9999. REM “frequency”: F (89, 1800) = 399.3, P<0.0001; “virus”: F (1, 1800) = 4.176e-009, P>0.9999. **F** left panel, episode number for 24h BL recordings divided by light (LP) and dark period (DP) and behavioural states. 2-Way ANOVA followed by Sidak’s *post-hoc* test using “state” and “virus” as factors (“state”: F (5, 126) = 232.1, P<0.0001; “virus”: F (1, 126) = 122.4, P<0.0001); centre panel, episode mean duration for each behavioural state during light and dark period. 2-Way ANOVA followed by Sidak’s *post-hoc* test using “state” and “virus” as factors (“state”: F (5, 126) = 60.36, P<0.0001; “virus”: F (1, 126) = 105.1, P<0.0001); right panel, ΔGluN1-LPO mice vigilance state transitions during light (top) and dark (bottom) period represented as percentage over GFP-LPO transitions. ΔGLUN1-LPO, *n* = 11; GFP-LPO, *n* = 12. In C, D, E and F data are represented as mean ± SEM. \**P* < 0.05, ** *P*< 0.005, *** *P*< 0.0005, † *P*< 0.00005 from multiple comparisons analysis corrected by Sidak’s (D, E and F) *post-hoc* test.

In addition to sleep loss and reduced cortical theta power, ΔGluN1-LPO mice had a highly fragmented sleep-wake phenotype: they lacked long wake and NREM sleep episodes, as they had significantly more wake and NREM sleep episodes (Figure 3F, left panel), with a decrease in their mean duration (Figure 3F, centre panel). Indeed, removing the NMDA receptor from LPO neurons increased transitions between wake and NREM sleep by > 55% (Figure 3F, right panel). The phenotype occurred in both the light and dark phases (Figure 3F). For REM sleep in ΔGluN1-LPO mice, there was a 50 to 70% decrease in episode number, mean duration and transitions to and from this vigilance state (Figure 3F).

### REGION-SPECIFIC EFFECT OF NMDA RECEPTOR ABLATION ON SLEEP-WAKE FRAGMENTATION

We also tested if deleting the GluN1 subunit in a region neighbouring the PO area, the anterior hypothalamic area (AHA), caused sleep loss or fragmentation (Supplementary Figure 4A-C). Bilateral injection o*f AAV-Cre-Venus* and *AAV-GFP* into the AHA of floxed-*grin1* mice, to generate ΔGluN1-AHA and GFP-AHA animals respectively, did not affect sleep and wake amounts during either the light or dark phases (Supplementary Figure 4A-C), nor did the deletion influence the number of transitions between vigilance states (Supplementary Figure 4D). ΔGluN1-AHA mice did not show any signs of sleep fragmentation: the episode number of NREM and REM sleep epochs and their mean duration were similar to GFP-AHA animals (Supplementary Figure 4E). Therefore, the fragmented sleep phenotype produced by deleting NMDA receptors originates region-selectively in the hypothalamus.

### THE INSOMNIA OF ΔNR1-LPO MICE PERSISTS UNDER HIGH SLEEP PRESSURE

To investigate if the fragmented sleep of ΔGluN1-LPO mice persisted under high sleep pressure and if NMDA receptors in LPO were required for sleep homeostasis, we performed 6h of sleep deprivation (SD) at the onset of the “lights on” period when the sleep drive is highest (Figure 4A). Although ΔNR1-LPO mice were awake and moving during the sleep deprivation, there were several indications that they were under high sleep pressure. During the sleep deprivation, the EEG theta power in ΔNR1-LPO mice was greatly reduced (red trace in Figure 4B when compared with GFP-LPO animals), and most of the power was concentrated in the delta frequency band (Figure 4B). Compared to GFP-LPO mice, ΔGluN1-LPO mice had many more sleep attempts during the sleep deprivation procedure (Figure 4C, left panel), and at the end of the 6 hour procedure they had a shorter latency to fall asleep (Figure 4C, right panel).

**Figure 4.**
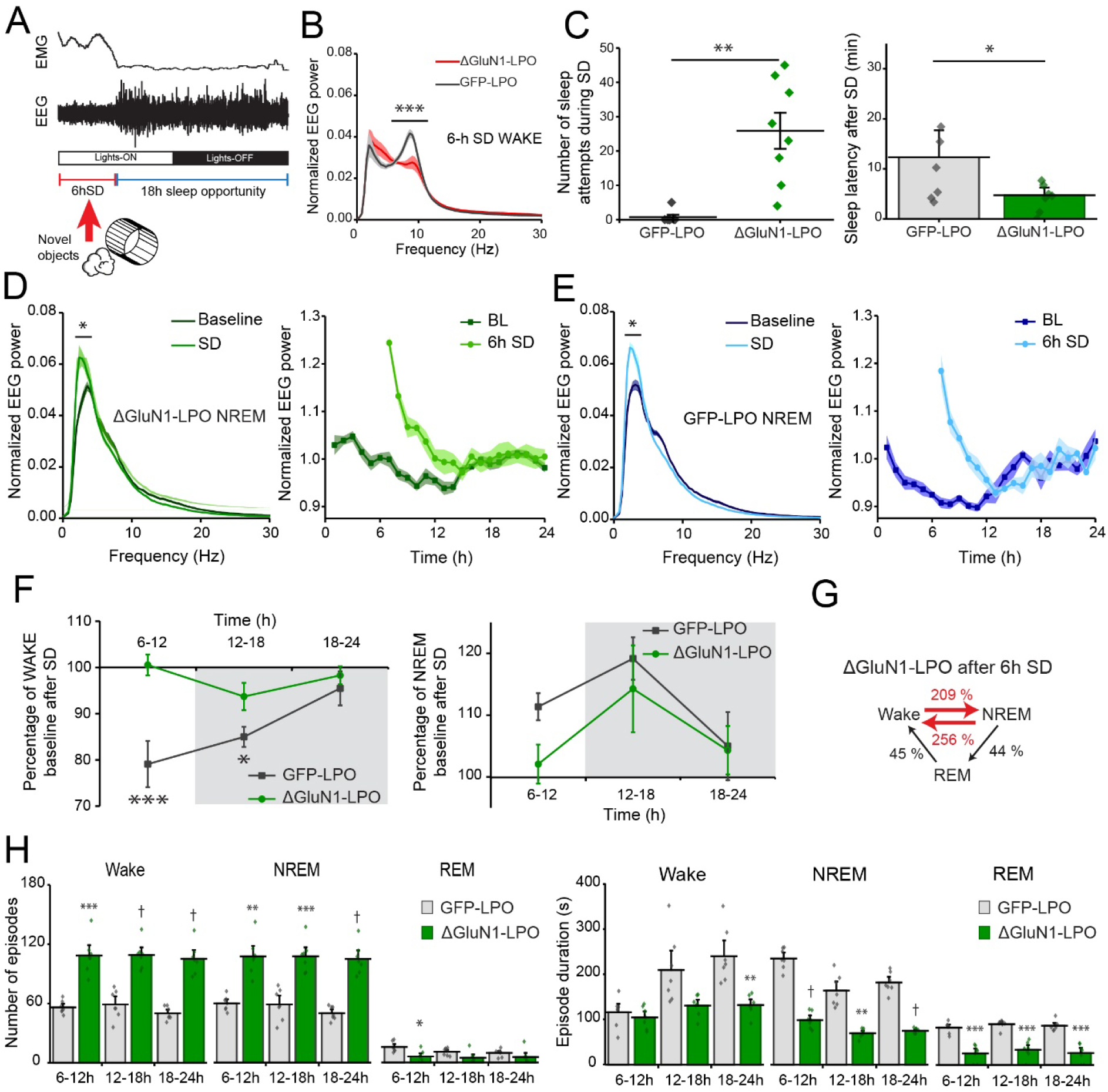
Deletion of the NMDA receptor in LPO does not affect sleep homeostasis and sleep-wake fragmentation still persists in the recovery sleep following sleep deprivation. **A**, representation of 6h SD protocol starting when lights turn ON, using novel objects to keep the animals awake. For the remaining 18h animals are left undisturbed. **B**, wake EEG power spectrum during 6h SD. 2-Way ANOVA followed by Sidak’s *post-hoc* test using “frequency” and “virus” as factors (“frequency”: F (89, 1080) = 125.1, P<0.0001; “virus”: F (1, 1080) = 4.318e-010, P>0.9999). **C**, number of sleep attempts during the 6h of SD (left) and latency to fall asleep (right) after SD, considering the first NREM bout as at least 30s long. Mann-Whitney test was used for sleep attempts (*P* = 0.0006) and unpaired Student’s *t* test for latency (*P* = 0.049). **D** and **E** left panels, NREM EEG power spectrum during 1h following 6h sleep deprivation (SD) compared with same circadian time during baseline recordings in ΔGluN1-LPO (D) and in GFP-LPO (E) animals. 2-Way ANOVA followed by Sidak’s *post-hoc* test using “frequency” and “virus” as factors. ΔGluN1-LPO “frequency”: F (89, 1080) = 653.1, P<0.0001; “virus”: F (1, 1080) = 8.999e-009, P>0.9999. GFP-LPO “frequency”: F (89, 1080) = 963.5, P<0.0001; “virus”: F (1, 1080) = 7.930e-009, P>0.9999; right panels, NREM EEG delta power calculated for every hour during baseline recordings and after 6h SD in ΔGluN1-LPO and GFP-LPO animals. The EEG power was normalized over the total power during each hour. **F**, percentage of wake (left) and NREM sleep amounts (right) in ΔGluN1-LPO and GFP-LPO mice over their own baseline after 6h SD. 2-Way ANOVA followed by Sidak’s *post-hoc* test using “time” and “virus” as factors (Wake “time”: F (1.564, 18.77) = 5.867, P=0.0151; “virus”: F (1, 12) = 10.69, P=0.0067. NREM “time”: F (1.479, 17.74) = 3.180, P=0.0782; “virus”: F (1, 12) = 2.906, P=0.1140). **G**, ΔGluN1-LPO transitions in the 18h following 6h SD presented as percentage of GFP-LPO mice transitions. **H** left panels, episode number calculated over 6h following SD for wake (left), NREM (centre) and REM sleep (right). 2-Way RM ANOVA followed by Sidak’s *post-hoc* test using “time” and “virus” as factors. Wake, “virus”: F (1, 12) = 79.90, P< 0.0001; “time”: F (1.852, 22.23) = 1.786, P= 0.1924. NREM, “virus”: F (1, 12) = 64.85, P< 0.0001, “time”: F (1.937, 23.24) = 1.780, P= 0.1916. REM, “virus”: F (1, 12) = 6.25, P= 0.0279, “time”: F (1.905, 22.87) = 6.702, P= 0.0056. Right panels, episode mean duration calculated by 6h following SD for wake (left), NREM (centre) and REM sleep (right). 2-Way RM ANOVA followed by Sidak’s *post-hoc* test using “time” and “virus” as factors. Wake, “virus”: F(1, 12) = 17.30, P= 0.0013, “time”: F (1.885, 22.62) = 13.10, P= 0.0002. NREM, “virus”: F (1, 12) = 202.5, P< 0.0001, “time”: F (1.676, 20.12) = 27.04, P < 0.0001. REM, “virus”: F (1, 12) = 63.84, P< 0.0001, “time”: F (1.853, 22.24) = 2.672, P= 0.0946. GFP-LPO, *n*=7; ΔGluN1-LPO, *n*=7. Data in all panels B, C, D, E, F and H are represented as means ± SEM. \**P* < 0.05, ** *P*< 0.005, *** *P*< 0.0005, † *P*< 0.00005 from multiple comparisons analysis using Sidak’s *post-hoc* correction.

During the subsequent first hour of sleep following sleep deprivation, both ΔGluN1-LPO and GFP-LPO mice had a significant increase in NREM delta power compared to their own baseline at the same circadian time (Figure 4D and E, left panels), showing that by this measure, sleep homeostasis was intact. The typical diurnal variation in EEG delta power over 24 hours seen in control mice was also still present in ΔGluN1-LPO mice (Figure 4D and E right panels). However, ΔGluN1-LPO mice were incapable of recuperating the sleep lost during sleep deprivation (Figure 4F). Following sleep deprivation as a percentage over their own baselines, GFP-LPO mice as expected increased their time asleep, reducing time spent awake. ΔGluN1-LPO animals, however, did not (Figure 4F). After sleep deprivation, ΔGluN1-LPO animals maintained a highly fragmented sleep phenotype during their recovery sleep, as shown by the over 2-fold increase over GFP-LPO mice in the number of transitions between NREM sleep and wake during the 18h sleep recovery opportunity time (Figure 4G). Additionally, NREM and wake episodes numbers were still increased, and mean duration decreased in ΔGluN1-LPO mice after SD, whereas REM sleep values were still consistently lower than in GFP-LPO mice (Figure 4H). The persistence of fragmentation after SD in ΔGluN1-LPO mice is particularly noteworthy, as under increased sleep pressure, quantified by the delta power rebound, sleep is deeper compared to baseline levels. These data suggest that ΔGluN1-LPO mice were extremely sleepy but could not stay asleep.

### SEDATIVES AND SLEEPING MEDICATION TRANSIENTLY IMPROVE SLEEP QUALITY OF ΔNR1-LPO MICE

We investigated whether drugs that induce NREM-like sleep, dexmedetomidine which is used in intensive care units for long-term sedation (Adams et al., 2013), and zolpidem (Ambien), a widely prescribed sleeping medication (Wisden et al., 2019), could reduce the high sleep fragmentation in ΔGluN1-LPO animals and restore consolidated sleep. Following *i.p*. injection of ΔGluN1-LPO animals, both dexmedetomidine (25 and 50 μg/kg) and zolpidem (5mg/kg) increased the time spent asleep (Figure 5A and B). 50 μg/kg of dexmedetomidine reduced the number of NREM sleep episodes and increased NREM episode mean duration for 1h after injection (Figure 3C left and right panels). Sleep-wake transitions were also reduced under both doses of dexmedetomidine (Figure 5D). On the other hand, zolpidem did not affect the number of NREM sleep episodes, but it increased their duration (Figure 5E, right and left panel), resulting in a decrease in transitions between behavioural states (Figure 5F). For both drugs, a few hours after administration the highly fragmented sleep pattern re-emerged (data not shown).

**Figure 5.**
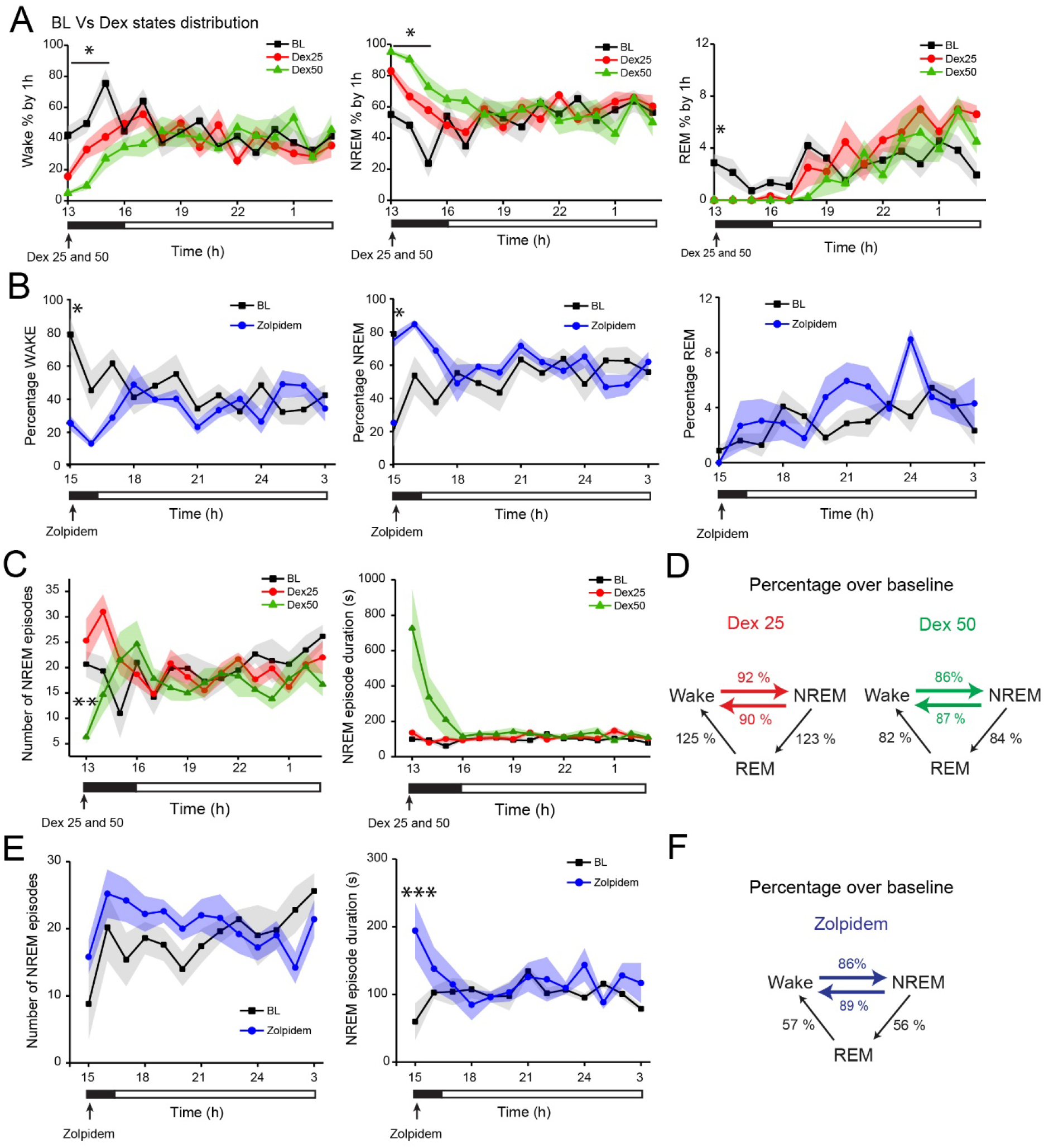
Dexmedetomidine and zolpidem both increase sleep time and dexmedetomidine reduces sleep fragmentation in mice lacking NMDA receptors in LPO. **A**, baseline states distribution as percentage of 1h in wake (left), NREM (centre) and REM sleep (right) compared with the distribution after injection of Dex 25μg/Kg (red) and Dex 50μg/Kg (green) at 13:00. 2-Way RM ANOVA followed by Tukey’s *post-hoc* test using “time” and “dose” as factors. For wake “time”: F (6.834, 102.5) = 3.244, *P*=0.0040; “dose”: F (2, 15) = 8.067, *P*=0.0042. For NREM “time”: F (6.768, 101.5) = 3.669, *P*=0.0016; “dose”: F (2, 15) = 7.578, *P*=0.0053. For REM “time”: F (5.394, 80.92) = 13.74, P<0.0001; “dose”: F (2, 15) = 1.808, *P*=0.1980). **B**, distribution by hour of wake (left), NREM (centre) and REM sleep (right) following injection of zolpidem 5mg/kg at 15:00. 2-Way RM ANOVA followed by Tukey’s *post-hoc* test using “time” and “dose” as factors. For wake, “time”: F (4.342, 34.73) = 1.596, P=0.1937; “dose”: F (1, 8) = 6.412, P=0.0351. For NREM sleep, “time”: F (4.377, 35.01) = 1.568, P=0.2006; “dose”: F (1, 8) = 6.006, P=0.0399. For REM sleep, “time”: F (4.315, 34.52) = 4.355, P=0.0050; “dose”: F (1, 8) = 2.192, P=0.1770. **C**, number of NREM episodes (left) and NREM episode mean duration (right) comparing baseline to dexmedetomidine (25 μg/kg and 50 μg/kg) sleep recordings. 2-Way RM ANOVA followed by Tukey’s *post-hoc* test using “time” and “dose” as factors (left panel: “time”: F (6.165, 92.48) = 1.850, *P*=0.1198; “dose”: F (2, 15) = 1.728, *P*=0.3142; right panel: “time”: F (1.266, 18.99) = 5.741, *P*=0.02098; “dose”: F (2, 15) = 6.516, *P*=0.0092). **D**, number of transitions during the “lights on” period after injection of dexmedetomidine 25 μg/kg (left) and 50 μg/kg (right), represented as percentage over the baseline recordings at the same circadian time. **E**, NREM episode number (left) and mean duration (right) comparing baseline to zolpidem (5 mg/kg) sleep recordings. 2-Way RM ANOVA followed by Tukey’s *post-hoc* test using “time” and “dose” as factors (left panel: “time”: F (5.530, 44.24) = 2.142, P=0.0720; “dose”: F (1, 8) = 0.6910, P=0.4299; right panel: “time”: F (12, 96) = 0.7856, P=0.6639; “dose”: F (1, 8) = 3.234, P=0.1098). **F**, number of transitions during the light period after injection of zolpidem represented as percentage over the baseline recordings at the same circadian time. For all panels, ΔGluN1-LPO, *n* = 6. Data in A, B, C and E are represented as mean ± SEM. \**P* < 0.05, ** *P*< 0.005 from multiple comparisons analysis using Tukey’s *post-hoc* correction.

### SLEEP FRAGMENTATION BUT NOT SLEEP LOSS IS PRODUCED BY SELECTIVE NMDA GluN1 SUBUNIT KNOCK-DOWN IN GABA LPO NEURONS

We identified an shRNA to selectively reduce GluN1 expression cell type selectively, for example, in GABAergic or glutamatergic cells in LPO. The efficacy of shRNAs to knockdown recombinant GluN1 cDNA expression was selected *in vitro* (Figure 6A and B). Having identified a suitable shRNA, *AAV-flex-shRNA-GluN1* and *AAV-flex-shRNA-scramble* were bilaterally injected into the LPO areas of Vgat-Cre and Vglut2-Cre mice (Figure 6C and Supplementary Figure 5A). Unlike the ΔGluN1-LPO mice, the overall macrostructure of vigilance states was not changed by GluN1 knockdown in GABA or glutamate neurons: Vgat-shRNA-GluN1 and Vglut2-shRNA-GluN1 mice did not have sleep loss compared with the respective scramble shRNA controls (Figure 6D and Supplementary Figure 5B). The EEG power spectra during wake, NREM and REM sleep were also not different, and there were no disruptions in REM sleep (Figure 6E and Supplementary Figure 5C). However, knock-down of the GluN1 subunit from LPO-Vgat neurons, but not LPO-Vglut2 neurons (Supplementary Figure 5D), caused a strong sleep-wake fragmentation phenotype (Figure 6F). For Vgat-shRNA-GluN1 mice, there were more episodes (Figure 6F, left panel), and decreases in episode duration (Figure 6F, centre panel), resembling the sleep fragmentation phenotype observed in ΔGluN1-LPO mice. There was a >55% increase in transitions between vigilance states in Vgat-shRNA-GluN1 animals compared with Vgat-shRNA-scr mice (Figure 6F, right panel), as ΔGluN1-LPO mice compared to their controls.

**Figure 6.**
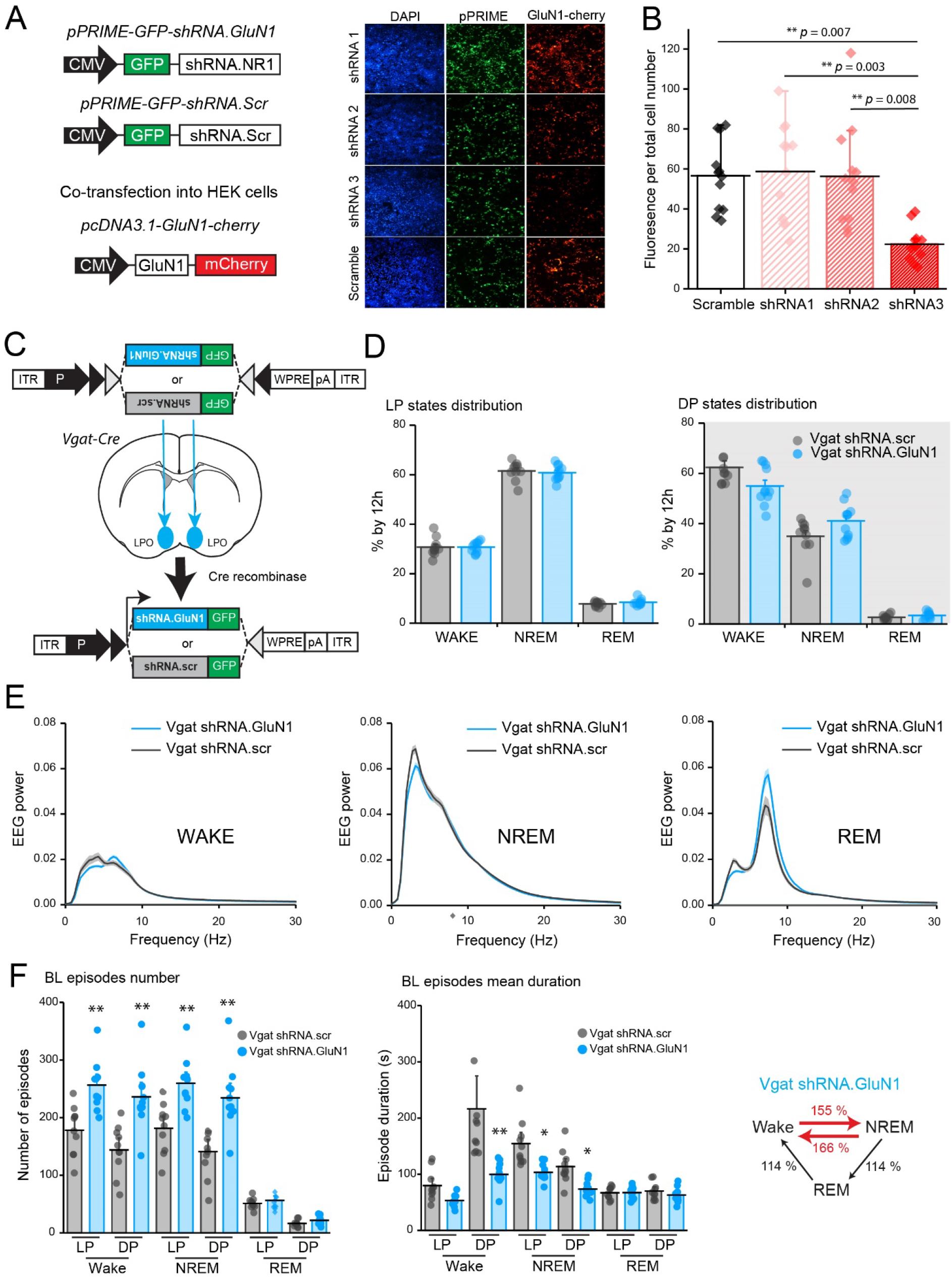
NMDA receptor GluN1 knock-down from GABA neurons in the LPO hypothalamus causes sleep-wake fragmentation but not sleep loss. **A**, DNA constructs used to test shRNA-GluN1 efficiency (left) and fluorescent images (right) of transfected HEK293 cells showing DAPI (left column, in blue), GFP reporter for expression of shRNA-GluN1 and shRNA-scramble pPRIME vectors (central column, in green), and mCherry reporter expression which indicates expression of the plasmid carrying the GluN1 sequence (right column, in red). **B**, quantification of mCherry fluorescence per total cell number after transfection of HEK293 cells with plasmids expressing the shRNA-GluN1 or the control scramble shRNA. *N* = 10, one-way ANOVA followed by Tukey’s *post-hoc* test for multiple comparisons (F = 9.126; *P*< 0.0001). **C**, *AAV-flex-shRNA-GluN1* or *AAV-flex-shRNA-scramble* (scr) were bilaterally injected in the LPO area of Vgat-cre animals to generate Vgat shRNA-GluN1 and Vgat-shRNA-scr animals. **D**, vigilance state distribution represented as % of 12h during light (left) and dark (right) periods for Vgat shRNA-GluN1 and Vgat-shRNA-scr animals. 2-Way ANOVA followed by Sidak’s *post-hoc* test using “state” and “virus” as factors (“state”: F (5, 113) = 563.6, P<0.0001; “virus”: F (1, 113) = 0.02612, P=0.8719). **E**, EEG power spectrum of wake (left), NREM (centre) and REM (right) sleep during 12h light period in Vgat-shRNA-GluN1 animals normalized over total EEG power. **F** left and centre panels, baseline episode number (left) and episode mean duration (centre) comparing Vgat-shRNA-GluN1 with control Vgat shRNA-scr mice. 2-Way ANOVA followed by Sidak’s *post-hoc* test using “state” and “virus” as factors. For episode number, “state”: F (5, 114) = 109.6, P<0.0001; “virus”: F (1, 114) = 72.99, P<0.0001. For mean duration, “state”: F (5, 114) = 39.01, P<0.0001; “virus”: F (1, 114) = 62.05, P<0.0001; right panel, number of transitions in Vgat shRNA-GluN1 mice represented as percentage over control group during 24h baseline recordings. Vgat-shRN-GluN1, *n* = 12; Vgat-shRNA-scr, *n* = 11. In B, D, E and F data are represented as mean ± SEM. In F, \**P* < 0.05, ** *P*< 0.005 from multiple comparison analysis corrected by Sidak’s *post-hoc* test.

As for the ΔGluN1-LPO mice, we next tested if the sleep-wake fragmentation phenotype of Vgat-shRNA-GluN1 mice persisted under conditions of strong sleep pressure following 6 hr of sleep deprivation. In contrast to pan-neuronal GluN1 deletion, Vgat shRNA-GluN1 animals did not show a “sleepy phenotype”, as their sleep attempts during sleep deprivation did not differ from the control group (Supplementary Figure 6A). Following 6hSD, Vgat-shRNA-GluN1 mice had a significant increase in the EEG delta power during the 1h following SD compared to their own baseline power, similarly to ΔNR1-LPO mice, showing that sleep homeostasis was intact (Supplementary Figure 6B). Vgat-shRNA-GluN1 mice maintained the fragmented sleep phenotype even under high sleep pressure, with more transitions (Supplementary Figure 6C), and more wake and NREM episodes (Supplementary Figure 6D) with decreased mean durations (Supplementary Figure 6E) compared with Vgat-shRNA-scr animals. Thus, the sleep-wake fragmentation aspect, but not the REM sleep loss, of ΔGluN1-LPO mice originates from GABA cells in LPO.

## DISCUSSION

The LPO hypothalamus is required for both NREM and REM sleep generation and NREM sleep homeostasis (Lu et al., 2002; Lu et al., 2000; Ma et al., 2019; McGinty and Sterman, 1968; Nauta, 1946; Reichert et al., 2019; Sherin et al., 1996; Szymusiak et al., 2007; Zhang et al., 2015). We explored how NMDA receptors on LPO neurons regulate sleep. We found that calcium levels in mouse LPO neurons, regardless of cell type, were highest and most sustained during REM sleep, although there was some “spikey activity” during NREM sleep. Deleting the core GluN1 subunit of NMDA receptors from LPO neurons substantially reduced the excitatory drive onto these cells and abolished calcium signals during all vigilance states. These ΔGluN1-LPO mice had less NREM sleep and no conventional REM sleep (atonia was present, but there was no EEG theta activation). In addition, ΔGluN1-LPO mice had highly fragmented sleep-wake: they had many more episodes of wake and NREM sleep, but each episode was shorter. Thus, AMPA glutamate receptor excitation alone on LPO sleep-promoting neurons is insufficient to maintain NREM sleep or produce REM sleep. The phenotype was further stratified. High sleep-wake fragmentation, but not sleep loss, was produced by selective GluN1 knock-down in GABAergic LPO neurons. The ΔGluN1-LPO mice phenotype is quite similar (wake-NREM fragmentation, loss of REM sleep) to mice with a double (global) deletion of the muscarinic receptor genes Chrm1 and Chrm3 (Niwa et al., 2018), so presumably this could intersect on the same pathway.

Our initial motivation for this study was testing the role of NMDA receptors in sleep homeostasis mediated by the PO hypothalamus. Calcium entry through NMDA receptors has been suggested to be part of the sleep homeostasis mechanism that tracks time spent awake (Liu et al., 2016). The sleep homeostasis process is reflected by changes in EEG delta power (Borbely et al., 2016). During the 24 hour cycle, delta power is highest during the “lights on” sleep phase and declines as each NREM sleep bout progresses (Figure 4D, E), which is thought to reflect the dissipation of the homeostatic sleep drive (Borbely et al., 2016). After sleep deprivation, neocortical activity in the subsequent (“recovery”) NREM sleep is deeper (more synchronised and thus has a higher delta power). However, we found that NMDA receptor deletion in LPO did not affect sleep homeostasis as defined by the classical criteria - EEG delta power showed its usual variation, increase and decrease over 24 hours. Even placing ΔGluN1-LPO mice under high sleep pressure by sleep deprivation did not enable the mice to sleep well. The sleep fragmentation persisted even during the recovery sleep, and the fragmented sleep started off with a higher delta power as expected for recovery sleep in the sleep homeostasis model. So, the sleep homeostatic process seems independent of the mechanism maintaining consolidated sleep. In fact, during sleep deprivation, ΔGluN1-LPO mice made multiple attempted entries to sleep. It is as if the mice were chronically sleepy, but they could not stay asleep.

Behavioural therapy is often ineffective for treating severe insomnia disorder, and medication remains an alternative approach if used cautiously (Shahid et al., 2012; Van Someren, 2020). Unlike sleep deprivation, which is usually efficient at inducing sleep, drugs could treat quite effectively the insomnia of ΔGluN1-LPO mice. Dexmedetomidine could transiently restore consolidated NREM sleep. Dexmedetomidine, an α2 adrenergic agonist, induces stage 3 NREM sleep in humans and NREM-like sleep in animals (Akeju et al., 2018; Gelegen et al., 2014; Zhang et al., 2015), and requires galanin/GABA neurons in LPO for its effects (Ma et al., 2019). Zolpidem (Ambien), a GABA_A_ receptor positive modulator, is a widely prescribed sleeping medication (Wisden et al., 2019). Its main effect in humans is to reduce latency to NREM sleep rather than maintaining consolidated sleep. Nevertheless, it did have an effect on ΔGluN1-LPO mice, restoring longer periods of NREM sleep, although not blocking the fragmentation.

Our findings demonstrate a novel aspect of REM sleep generation. REM sleep is characterized by a high theta delta ratio in the EEG and muscle atonia. In rodents, the theta itself detected in the cortical EEG seems to originate mostly from the hippocampus. Indeed, the theta activation during REM is required for memory processing (Boyce et al., 2016; Izawa et al., 2019). Although the brainstem circuitry that generates muscle atonia during REM sleep is reasonably well understood, the circuitry that produces the wake-like activation (theta activity in the EEG) of the neocortex during REM sleep is only partially characterized, seeming to require distributed circuitry throughout the forebrain (Izawa et al., 2019; Luppi et al., 2017; Peever and Fuller, 2016; Renouard et al., 2015; Yamada and Ueda, 2019), including the MCH NREM-REM promoting neurons in the lateral hypothalamus (Jego et al., 2013), and REM-off and REM-on neurons in the dorsal medial hypothalamus (Chen et al., 2018), and GABA and cholinergic neurons in the medial septum which project to the hippocampus (Yoder and Pang, 2005). Although LPO has long been known to be required for REM sleep (Lu et al., 2002; Lu et al., 2000), we were surprised to discover that LPO neurons, regardless of type (e.g. galanin, Vgat, Vglut2), have actually their highest calcium activity during REM sleep. We found that GluN1 knockdown in GABA cells of LPO did not influence REM sleep, whereas the pan knockout in all LPO neurons did, so the cell type(s) responsible for REM sleep generation in LPO require further investigation.

NMDA receptor properties could be responsible for maintaining NREM and REM sleep promoting LPO neurons in the “on” state. As well as calcium permeability, NMDA receptors have a voltage-dependent magnesium block, and in contrast to AMPA-gated ionotropic glutamate receptors, with which they are often paired in synapses, stay open from around 100 msec to 1 s (Paoletti et al., 2013). Because of these properties NMDA receptors are famously studied for their role in synaptic plasticity, and this is the thinking behind why NMDA receptors would be involved in sleep homeostasis (Liu et al., 2016; Raccuglia et al., 2019). But these same properties also allow NMDA receptors to act as pacemakers, controlling rhythmic firing e.g. in those circuits involved in breathing, swimming and walking (Li et al., 2010; Steenland et al., 2008). It is also possible that the long open times of NMDA receptors stabilize the sleep-on neurons in their firing mode. It will be interesting to see if this role of NMDA receptors generalizes to other sleep-promoting circuits. For example, we previously found that genetic silencing of mouse lateral habenula neurons with tetanus toxin light chain produced high NREM sleep-wake fragmentation with conserved amounts of total sleep and wake, with the fragmentation effect mostly occurring during the “lights-on” sleep phase (Gelegen et al., 2018). It seems likely that disrupting NMDA receptors on these cells would also produce insomnia, given that that NMDA receptors are needed to keep lateral habenula cells in burst firing (active) mode (Cui et al., 2019; Yang et al., 2018).

In conclusion, we have found that cells in the LPO hypothalamus have their highest calcium activity during REM sleep. This calcium is determined by NMDA receptors. Reducing NMDA receptors in the LPO hypothalamic area causes profound insomnia (wake-NREM sleep fragmentation) and substantial loss of theta activity during REM sleep. Thus, glutamate activation, via NMDA receptors, is essential for NREM sleep maintenance, and drives a key characteristic of REM sleep (theta power).

## Supporting information

Miracca.Supp figures and legends

## ACKNOWLEDGEMENTS

Funded by the Wellcome Trust (107839/Z/15/Z, N.P.F. and 107841/Z/15/Z, W.W.), the UK Dementia Research Institute (N.P.F. and W.W.), and an Imperial College Schrödinger Scholarship (G.M.).

## MATERIAL AND METHODS

### Mice

All experiments were performed in accordance with the United Kingdom Home Office Animal Procedures Act (1986) and were approved by the Imperial College Ethical Review Committee. Wild-type C57BL/6N mice were purchased from Charles River at 7/8 weeks of age. Grin1^flox^ (also known as NR1^flox^ or fNR1) mice (Tsien et al., 1996) were purchased from The Jackson Laboratory (Jack stock number 005246) after kind donation by S. Tonegawa. Vglut2-Cre animals (*Vglut2-ires-Cre: Slc17a6^tm2(cre)Lowl^/J*) and Vgat-Cre mice (*Vgat-ires-Cre: Slc32a1^tm2(cre)Lowl^/J*) were kindly provided by B.B. Lowell and purchased from The Jackson Laboratory (JAX stock 016963 and 016962)(Vong et al., 2011). Galanin-Cre mice (Tg(Gal-cre)KI87Gsat/Mmucd) were generated by GENSAT and kindly deposited at the Mutant Mouse Regional Resource Center, stock No. 031060-UCD, GENSAT-Project (NINDS Contracts N01NS02331 & HHSN271200723701C to The Rockefeller University, New York) (Schmidt et al., 2013). Nos-Cre animals (*Nos1-ires-Cre^tm1(cre)Mgmj^/J*), were kindly provided by M. G. Myers, and purchased from The Jackson Laboratory (JAX stock 017526) (Leshan et al., 2012).

All mice were housed at a maximum of five mice per cage with food and water *ad libitum* and maintained under the same conditions (21±1C, reversed 12h dark/light cycle starting at 4:00 AM). For behavioural experiments, mice were singly housed, and experiments performed during lights-OFF unless otherwise specified, while photometry recordings were performed during lights-ON.

### Transgenes and AAVs

All AAVs (serotype 1/2) were produced in house. The adenovirus helper plasmid pFΔ6, the AAV helper plasmids pH21 (AAV1) and pRVI (AAV2), and the pAAV transgene plasmids were co-transfected into HEK293 cells and the resulting AAVs collected on heparin columns, as described previously (Klugmann et al., 2005; Yu et al., 2015). Plasmid *pAAV-iCre-2A-Venus* was provided by Thomas Kuner (Abraham et al., 2010). Plasmid *pAAV-GFP* was a gift from John T. Gray (Addgene plasmid 32396). To create *pAAV-hsyn-GCaMP6s* and the *pAAV-hsyn-flex-GCaMP6s*, the *GCaMP6s* reading frame from *pGP-CMV-GCaMP6s* (Addgene plasmid 40753, gift of Douglas Kim) (Chen et al., 2013) was mutated into *pAAV-flex-hM3Dq-mCHERRY (Krashes et al., 2011*), either removing the *flex-M3-cherry* component or keeping both sets of *loxP* sites (*pAAV-flex* backbone), respectively. For knocking down GluN1 expression cell type-selectively, the pPRIME system (Stegmeier et al., 2005), cloned into AAV transgenes, was used to generate shRNAs – see section (“Generation of shRNAs to target GluN1”).

### Surgeries

All surgeries used adult male and female mice, 8-12 weeks old and were performed under deep general anaesthesia with isoflurane (3% induction/ 2% maintenance) and under sterile conditions. Before starting the surgery, mice were injected subcutaneously (s.c.) with Buprenorphine (Vetergesic 0.3 mg/mL, 1:20 dilution in 0.9% sterile saline solution, final 0.1mg/kg) and Carprofen (Rimadyl 50mg/mL, 1:50 dilution in 0.9% sterile saline solution, final 5 mg/kg) and then placed in a stereotaxic frame. Mouse core temperature was constantly checked by rectal probe while respiration rate was regularly checked by eye.

For AAV injections, the virus was injected at a rate of 0.1 μL/min using Hamilton microliter #701 10 μL syringes and a stainless-steel needle (33-gauge, 15 mm long). LPO coordinates used for bilateral injection sites were relative to Bregma: AP, +0.40; ML, -/+ 0.75, DV was consecutive, injecting half volume at +5.20 and half at +5.15. A total volume of 0.3 μL each side was injected. Control AHA coordinates used for bilateral injections sites were relative to Bregma: AP, −0.58; ML, -/+ 0.65; DV, +5.60 and +5.50 for consecutive injections. Mice injected with AAVs were allowed 1 month for recovering and for the viral transgenes to adequately express before being fitted with Neurologger 2A devices (see below) and undergoing any experimental procedures.

For sleep recordings, EEG screw electrodes were chronically implanted on mice skull and EMG wire electrodes (AS634, Coorner Wire) were inserted in the neck extensor muscles. EEG screws were placed on the skull at: −1.5 mm midline, +1.5 mm Bregma; −1.5 mm midline, −2 mm Bregma; +1.5 mm midline, −2 mm Bregma.

For fibre photometry, a monofibre (Ø 200 μm, 0.37 NA, Doric Lenses) was chronically implanted together with EEG and EMG electrodes. The fibre was positioned after AAV injections above the LPO following coordinates relative to Bregma: AP, +0.10; ML, - 0.90, DV, −5.00 mm.

For all surgeries, the wound was sewed around the headstage and the mouse was left recovering in a heat box. All instrumented mice were single housed to avoid lesions to the headstage.

### EEG/EMG recordings and analysis

EEG and EMG traces were recorded using Neurologger 2A devices as described before (Anisimov et al., 2014; Gelegen et al., 2014; Vyssotski et al., 2009), at a sampling rate of 200 Hz. The data obtained from the Neurologger 2A were downloaded and visualized using Spike2 Software (Cambridge Electronic Design, Cambridge, UK). The EEG was high pass filtered (0.5Hz, −3dB) using a digital filter, while EMG was band pass filtered between 5-45 Hz (−3dB). To define the vigilance states of Wake, NREM and REM sleep, delta power (0.5-4.5 Hz) and theta ratio (theta power [5-10 Hz]/delta power) were calculated, as well as the EMG integral. Automated sleep scoring was performed using a Spike2 script and the result was manually corrected. For the three vigilance states, amounts percentages were calculate using costume Spike2 scripts. For sleep architecture analysis, costume MATLAB scripts were used. Fast Fourier transformation (512 points) was used to calculate EEG power spectra.

### Sleep deprivation protocol and drug testing

Mice were fitted with Neurologger 2A devices and the 24h sleep-wake baseline (BL) was recorded. After BL, mice with Neurologger 2A devices were sleep deprived from the light period onset (4:00pm) for 6 hours, introducing novel objects in their home cage (Tobler et al., 1997). To make the procedure minimally stressful, mice were never touched, apart from when changing cages. Sleep recordings were stopped at the end of the dark period of the following day (4:00pm).

Dexmedetomidine injections were prepared from stock solution of 0.5 mg/mL (Dexdomitor), diluted in sterile saline before injections. Mice were *i.p*. injected with the dose of 25 or 50 μg/kg at 1:00 pm (lights-OFF) to record sleep and fragmentation phenotype. Zolpidem injections were prepared by dissolving zolpidem tartrate powder (Sigma-Aldrich) in sterile saline. Mice were injected *i.p*. with 5 mg/kg of solution at 3:00 pm (light-OFF) and their EEG/EMG traces recorded for at least 24 hours.

### Histology and immunostaining

Animals were perfused transcardially with 20 mL of cold 1x PBS at a rate on 4 mL/min, followed by 20 mL 4% paraformaldehyde (PFA, 4mL/min) in 1x PBS. Brains were dissected and post-fixed in 4% PFA overnight, and then transferred in 30% sucrose. After 3 days in sucrose, brains were cut in 35-μm coronal slice using a microtome (Leica). For staining, slices were transferred in an epitope retrieval solution (0.05% Tween-20, 10 mM sodium citrate buffer, and pH 6.0) for 20min at 82°C, then left at room temperature (RT) for 15min before being washed. After 3 washes of 10min in 1x PBS, brain slices were blocked in 0.2% Triton™ x-100 (Sigma-Aldrich), 20% Normal Goat Serum (NGS, Vector Laboratories) in 1x PBS for 1h at RT, shaking. Primary antibody staining was then performed overnight at 4°C shaking in 0.2% Triton, 2% NGS in 1x PBS. In case of double staining, both primary antibodies were added in the solution unless cross-reactivity was previously observed. The following day, slices were washed 3 times in 1x PBS for 10 minutes and then secondary antibody solution was applied for 1h and 30 min in 0.2% Triton, 2% NGS in 1x PBS, at RT shacking. If double staining was required, washes and another secondary antibody incubation were carried out. For anti-IBA1 staining, donkey normal serum was used instead that NGS with same dilutions. After secondary antibodies incubations, slices were washed again for 3 times for 10min, RT in 1x PBS shacking and DAPI staining (1:5000 in PBS, Hoechst 33342, Life Technologies) was then performed for a maximum of 10min. After at least 1 wash in 1x PBS, slices were ready to be mounted. For mounting, microscope slides (Superfrost PLUS, Thermo Scientific), mounting media ProLong™ Gold Antifade Reagent (Invitrogen) and glass cover slides (24 x 50 mm, VWR Internartional) were used. Primary antibodies: rabbit anti-GFP (Invitrogen, A6455, 1:1000), chicken anti-GFP (Abcam, ab13970, 1:1000), rabbit anti-GFAP (Dako, Z0334, 1:500), goat anti-IBA-1 (Abcam, ab5076, 1:500), rat anti-mCherry (Invitrogen, M11217, 1:1000). Secondary antibodies (all from Invitrogen): Alexa Fluor-488 goat anti-Chicken (A11039, 1:500), Alexa Fluor-488 goat anti-rabbit (A11008, 1:500), Alexa Fluor-594 goat anti rabbit (A11072, 1:500) and Alexa Fluor-594 donkey anti-goat (A11058, 1:500), Alexa Fluor-568 goat anti-rat (A11077, 1:500).

### Acute slice preparation and electrophysiology recordings

Mice were euthanized by cervical dislocation and subsequent decapitation. The brain was rapidly retrieved to be sliced and placed into cold oxygenated N-Methyl-D-glucamine (NMDG) solution (in mM: NMDG 93, HCl 93, KCl 2.5, NaH2PO4 1.2, NaHCO3 30, HEPES 20, glucose 25, sodium ascorbate 5, Thiourea 2, sodium pyruvate 3, MgSO4 10, CaCl2 0.5). Para-horizontal slices (thickness 300 μm) encompassing the LPO area were obtained using a vibrotome (Vibrating Microtome 7000smz-2; Campden Instruments LTD, UK). Slices were incubated for 15min in NMDG solution at 33 °C with constant oxygenation, and transferred to oxygenated standard aCSF (in mM: NaCl 120, KCl 3.5, NaH2PO4 1.25, NaHCO3 25, glucose 10, MgCl2 1, CaCl2 2) solution for at least 1 hour at room temperature. Slices were transferred to a submersion recording chamber and were continuously perfused at a rate of 4-5ml/min with fully oxygenated aCSF at room temperature. For whole-cell recording, patch pipettes at 4-6 MΩ were pulled from borosilicate glass capillaries (1.5mm OD, 0.86 mm ID, Harvard Apparatus, #GC150F-10) and filled with intracellular solution containing (in mM: 128 CsCH3SO3, 2.8 NaCl, 20 HEPES, 0.4 EGTA, 5 TEA-Cl, 2 Mg-ATP, 0.5 NaGTP (pH 7.35, osmolality 285mOsm). 0.1% Neurobiotin was included in the intracellular solutions to identify the cell position and morphology following recording. Recordings were performed using a Multiclamp 700B amplifier (Molecular Devices. CA). Access and input resistances were monitored throughout the experiments. The access resistance was typically < 20 MΩ, and results we discarded if resistance changed by more than 20%.

GFP+ neurons were visually identified and randomly selected. For AMPA and NMDA current, a bipolar stimulus microelectrode (MX21AEW, FHC) was placed 100-200um away from recording site caudally. The intensity of stimulus (10ms) was adjusted to evoke a measurable, evoked EPSC in recording cells. AMPA and NMDA mixed currents were measured at a holding potential of +40mV. After obtaining at least 10 sweeps of stable mixed currents, D-AP5 (50 μM) was perfused to bath solution for 15min and AMPA currents were measured. NMDA currents were obtained by subtracting AMPA currents from mixed currents off-line. The peak amplitude of both currents was used for AMDA/NMDA ratio analysis. For sEPSCs, GFP+ LPO neurons were voltage clamped at −70mV constantly. A stable baseline recording was obtained for 5-10 min. Frequency, amplitude, rise & decay time constants of sEPSCs were analysed off-line with the Mini Analysis (Synaptosoft). Frequency was obtained from 2 min of recording. All recordings were made under the presence of picrotoxin (100 μM).

For immunohistochemistry following electrophysiological recordings, brain slices were post-fixed in 4% PFA overnight at 4 °C. PFA was then washed away 3 times for 10min in 1x PBS and slices were blocked and permeabilized in 20% NGS or 2% Bovine Serum Albumin (BSA) for 3 hours shaking. Primary anti-GFP Ab to trace viral distribution was diluted in 2% NGS, 0.5/0.7% TritonX in PBS overnight at 4°C shaking. After 4 washes in 1x PBS fpr 10min each, secondary Ab was diluted in 2% NGS and 0.5% TritonX for 3 hours at RT and shaking. After washes and to track Neruobiotin filled neurons recorded by electrophysiology, an Alexa594-conjugated streptavidin (Invitrogen) was diluted 1:500 in 1% NGS, 0.5% TritonX and slices were incubated for 2-3 hours at RT. 4 washes of 15min and subsequent DAPI incubation for 10min were performed before slices were mounted on glass slides as described in Section 2.8.

### Calcium Photometry

Following 4 weeks of recovery, mice were acclimatized to the testing environment for at least 2 hours before behavioural experiments and then recorded for 6 hours during the light period. As light source, a Grass SD9 stimulator was used to control a 473 nm Diode-pumped solid state (DPSS) laser with fibre coupler (Shanghai Laser & Optics century Co.) and adjustable power supply (Shanghai Laser & Optics century Co.). A lock-in amplifier (SR810, Stanford Research Systems, California, USA) was used to drive the laser at 125 Hz TTL pulsations with an average power of 80 μW at the tip of the fibre directly connected to the mouse. The light source was joined to a fluorescence cube (FMC_GFP_FC, Doric Lenses) through an optical fibre patch cord (Ø 200 μm, 0.22 NA, Doric Lenses). From the filter cube, an optical patch cords (Ø 200 μm, 0.37 NA, Doric Lenses) was connected to the monofibre chronically implanted in the mouse brain using a ceramic sleeves (Thorlabs). The GCaMP6s output was then filtered at 500-550 nm through the fluorescence cube, converted in Volts by a photodiode (APD-FC, Doric Lenses) and then amplified by the lock-in amplifier with a time constant of 30ms. Finally, the signal was digitalized using a CED 1401 Micro box (Cambridge Electronic Design, Cambridge, UK) and recorded at 200 Hz using Spike2 software (Cambridge Electronic Design, Cambridge, UK). Photometry, EEG and EMG data were aligned offline using Spike2 software and analyzed using custom made MATLAB (MathWorks) scripts. For each experiment, the photometry signal *F* was normalized to baseline using the function Δ*F/F* = (*F-F_0_*)/*F_0_*, where *F*_0_ is the mean fluorescence across the signal analyzed.

### Generation of shRNAs to target GluN1

We used the <u>p</u>otent <u>R</u>NA <u>i</u>nterference using <u>m</u>icroRNA <u>e</u>xpression (PRIME) system, where shRNAs are placed into the context of mir30 microRNA sequence (Stegmeier et al., 2005). By consulting the website *http://katahdin.cshl.org*, three shRNAs were designed to target exons 11 to 18 of the *Grin1* gene, encoding the region from amino acids 409 to 683 of GluN1. The sequences were amplified from mouse genomic DNA using primers (pSM2C Forward: 5’-GATGGCTG-CTCGAG-AAGGTATAT-TGCTGTTGACAGTGAGCG-3’; pSM2C Reverse: 5’-GTCTAGAG-GAATTC-CGAGGCAGTAGGCA-3’), following the protocol previously described (Stegmeier et al., 2005).

The three sequences were referred as shRNA-GluN1-1.1, −2 or −3 (the underlined sequences are the 22mers specific for the GluN1 subunit):

shRNA-GluN1.1 targeted the GluN1 sequence at 1800bp (600aa) of the CDS: 3’-TGCTGTTGACAGTGAGCG<u>AACTGACCCTGTCCTCTGCCAT</u>TAGTGAAGCCACAGATGTA<u>ATGGCAGAGGACAGGGTCAGTG</u>TGCCTACTGCCTCGGA-5’
shRNA-GluN1.2 matched the GluN1 sequence from 2565bp (855aa) of the CDS: 3’-TGCTGTTGACAGTGAG<u>CGCGCCGTGAACGTGTGGAGGAAG</u>TAGTGAAGCCACAGATGTA<u>CTTCCTCCACACGTTCACGGCT</u>TGCCTACTGCCTCGGA-5’
shRNA-GlN1.3 targeted the GluN1 sequence at 2215bp (738aa) of the CDS: 3’-TGCTGTTGACAGTGAGCG<u>CGGAGTTTGAGGCTTCACAGAA</u>TAGTGAAGCCACAGATGT<u>ATTCTGTGAAGCCTCAAACTCCA</u>TGCCTACTGCCTCGGA-5’

As control for shRNA-GluN1 sequences, an shRNA scramble hairpin was also produced, making sure it would not be complementary to any sequences in the mouse DNA. The shRNA-scramble sequence was:

3’-GCTGTTGACAGTGAGCGAGCTCCCTGAATTGGAATCCTAGTGAAGCCACAGATGTAGGATTCCAATTCAGCGGGAGCCTGCCTACTGCCTCGGA-5’.

The three shRNA-GluN1 hairpins and the shRNA-scramble hairpin were cloned into the pPRIME vector in the context of the mir30 micro RNA sequence, to be then expressed and tested in HEK293 cells. To establish shRNA efficiencies in knocking down the NMDA GluN1 subunit expression, a plasmid was constructed expressing GluN1-2A-mCherry under the control of the CMV promoter. Each GluN1 shRNA pPRIME plasmid was then transfected into HEK293 cells together with pGluN1-2A-mCherry. After 60 hours in culture, cherry red fluorescence was quantified. The GluN1.3 shRNA produced lower fluorescence intensity, and thus higher inhibition of GluN1 expression, and it was therefore cloned into an AAV transgene in an inverse orientation flanked by lox sites, as we described previously (Yu et al., 2015), to produce *AAV-flex-shRNA-GluN1*. The transgene expresses GFP as well as shRNA-GluN1.

### STATISTICAL ANALYSIS

Origin, MATLAB and GraphPad Prism 8 ware used for graphs and statistical analysis. Data collection and experimental procedure conditions were randomized. The experimenter was not blinded during the procedures. Data are presented as mean ± standard error of the means (SEM). Normality of each data set distribution was tested using the Kolmogorov-Smirnoff test. Unpaired two-tailed Student’s *t* test or one-way ANOVA were used to compare groups when only one variable was present. For measurement and data collected over time or with two separate independent variables, a 2-way repeated measures (RM) ANOVA or a simple 2-way ANOVA followed by *post-hoc* Sidak or Tukey tests were performed. When data did not result as normally distributed, Mann-Whitney test was performed. All details of statistical analyses used are specified in each figure legend. Statistical significance was considered when \**P* < 0.05.

