## Supplementary material for "Hypothalamic NMDA receptors stabilize NREM sleep and are essential for REM sleep": Miracca.Supp figures and legends

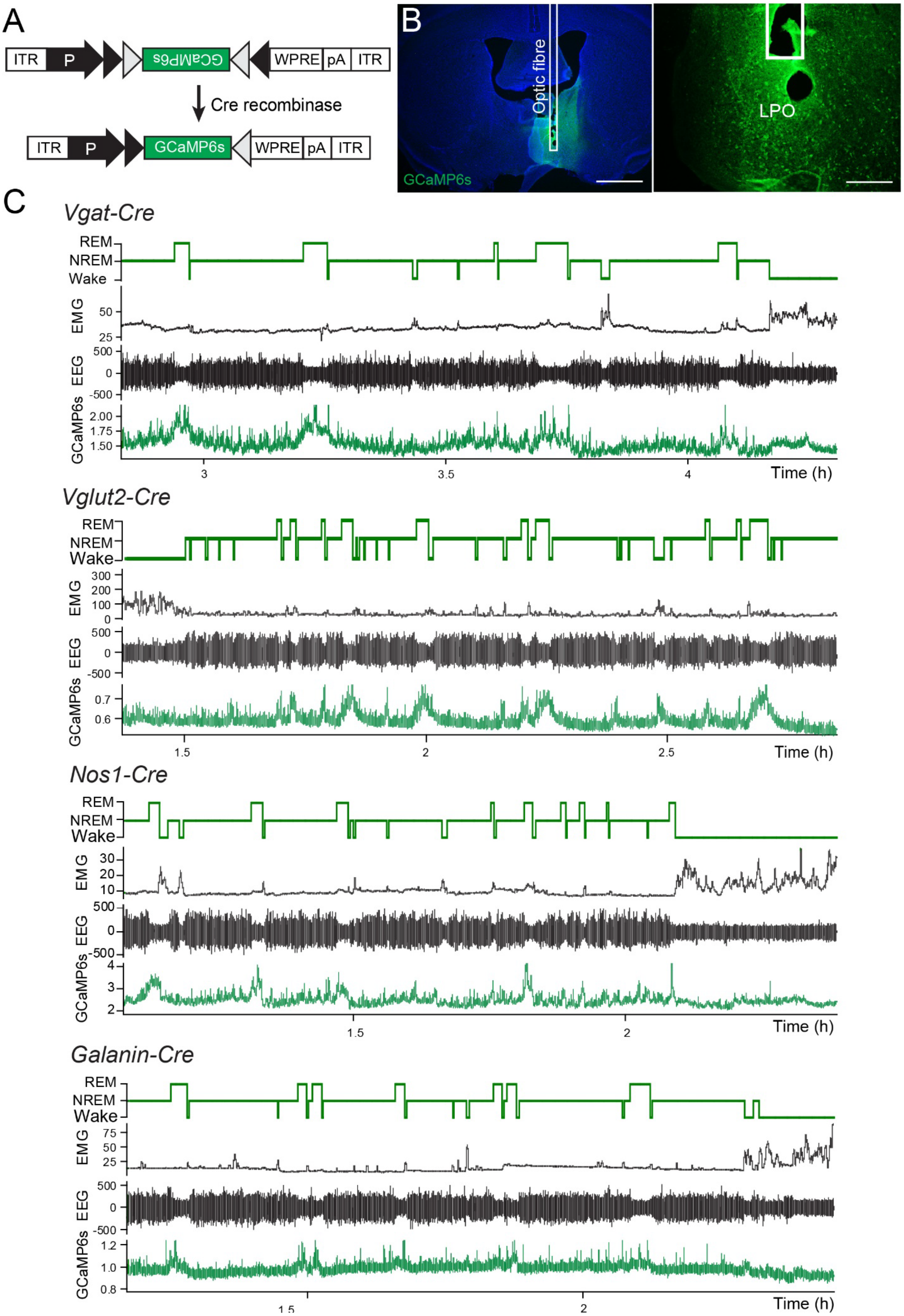

A
ITR
P
GCaMP6s
WPRE
pA
ITR
Cre recombinase
ITR
P
GCaMP6s
WPRE
pA
ITR
B
Optic fibre
GCaMP6s
LPO
C
Vgat-Cre
Vglut2-Cre
Nos1-Cre
Galanin-Cre
REM
NREM
Wake
EMG
EEG
GCaMP6s
Time (h)

**Supplementary Figure 1. LPO neurons are most active during REM sleep.**

**Supports Figure 1.**

**A**, *AAV-flex-GCaMP6s* transgene used in Cre-expressing mouse lines. **B**, immunohistochemistry from a Vgat-Cre mouse showing the fibre photometry track ending in the LPO area (right) and expression of the GCaMP6s calcium sensor (left; green = GFP/GCaMP6s, blue = DAPI). Scale bar, 1 mm on the left and 200 μm on the right. **C**, examples of photometry traces in Vgat-Cre, Vglut2-Cre, Nos1-Cre and Galanin-Cre mice. Each example shows from the top: hypnogram, EMG, EEG and GCaMP6s photometry traces.

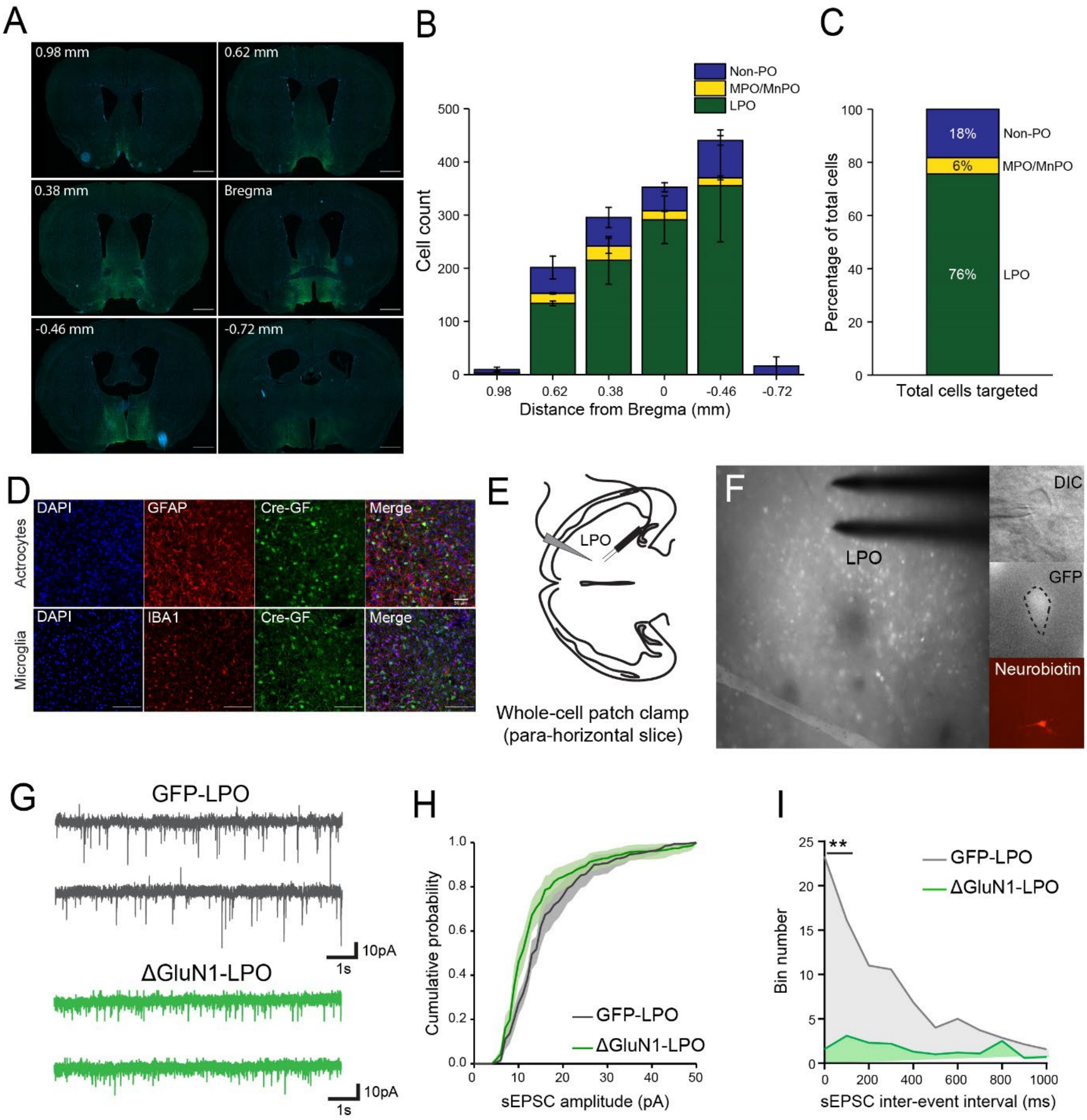
A
0.98 mm
0.62 mm
0.38 mm
Bregma
-0.46 mm
-0.72 mm
B
Non-PO
MPO/MnPO
LPO
Cell count
Distance from Bregma (mm)
C
18%
6%
76%
Non-PO
MPO/MnPO
LPO
Percentage of total cells
Total cells targeted
D
DAPI
GFAP
Cre-GF
Merge
IBA1
Actrocytes
Microglia
E
LPO
Whole-cell patch clamp
(para-horizontal slice)
F
LPO
DIC
GFP
Neurobiotin
G
GFP-LPO
ΔGluN1-LPO
10pA
1s
H
Cumulative probability
sEPSC amplitude (pA)
GFP-LPO
ΔGluN1-LPO
I
**
GFP-LPO
ΔGluN1-LPO
Bin number
sEPSC inter-event interval (ms)

**Supplementary Figure 2.** ***AAV-Cre-Venus*** **mapping and characterization of spontaneous excitatory post-synaptic currents (sEPSCs) in ΔGluN1-LPO mice**.

**Supports Figure 2.**

**A**, mapping of Cre recombinase expression using immunohistochemistry (Cre-venus in green; DAPI in blue). Coordinates are relative to Bregma. Scale bars, 1mm. **B**, distribution of *AAV-Cre-2A-Venus* virus based on distance from Bregma quantified by counting green (Venus-positive) cell bodies. “Non-PO” refers to areas not part of the larger preoptic structure; MPO/MnPO, medial and median preoptic. **C**, percentage of cells expressing Cre recombinase based on the anatomical structure. **D**, double-label immunohistochemistry investigating if the *AAV-Cre-2A* transgene expressed in astrocytes or microglia. From left to right: DAPI (blue), glial fibrillary acidic protein (GFAP) for astrocytes (red, top row) and ionized calcium binding adaptor molecule 1 (IBA1) for microglia (red, bottom raw), Cre recombinase (green) and merged images. Scale bars, 100 µm. **E**, schematic of para-horizontal slicing used to record evokes and spontaneous EPSCs from LPO neurons. **F**, microscope images of GFP+ (Venus-positive) cells transduced with *AAV-Cre-2A-Venus* in LPO. The cell successfully patched is shown in differential interface contrast (DIC), grey-scale for GFP and neurobiotin immune-detection. **G**, example of sEPSC recordings in GFP-LPO (top, in grey) and ΔGluN1-LPO (bottom, in green) neurons. **H**, Cumulative probability histogram of sEPSCs amplitude. 2-way ANOVA followed by Sidak’s *post-hoc* test using “probability” and “virus” as factors (“probability”: F (50, 765) = 129.8, P<0.0001; “virus”: F (1, 765) = 25.24, P<0.0001). **I**, cumulative bin number of inter-interval events (IEI) of sEPSCs. 2-way ANOVA followed by Sidak’s *post-hoc* test using “probability” and “virus” as factors (“probability”: F (25, 390) = 3.068, P<0.0001; “virus”: F (1, 390) = 15.31, P=0.0001). For B and C, *n*=3; for H and I, GFP-LPO, *n*=7; ΔGluN1-LPO, *n*=10. In B, C and H data are represented as mean ± SEM. In I, **P < 0.005 from multiple comparison analysis corrected by Sidak’s *post-hoc* test.

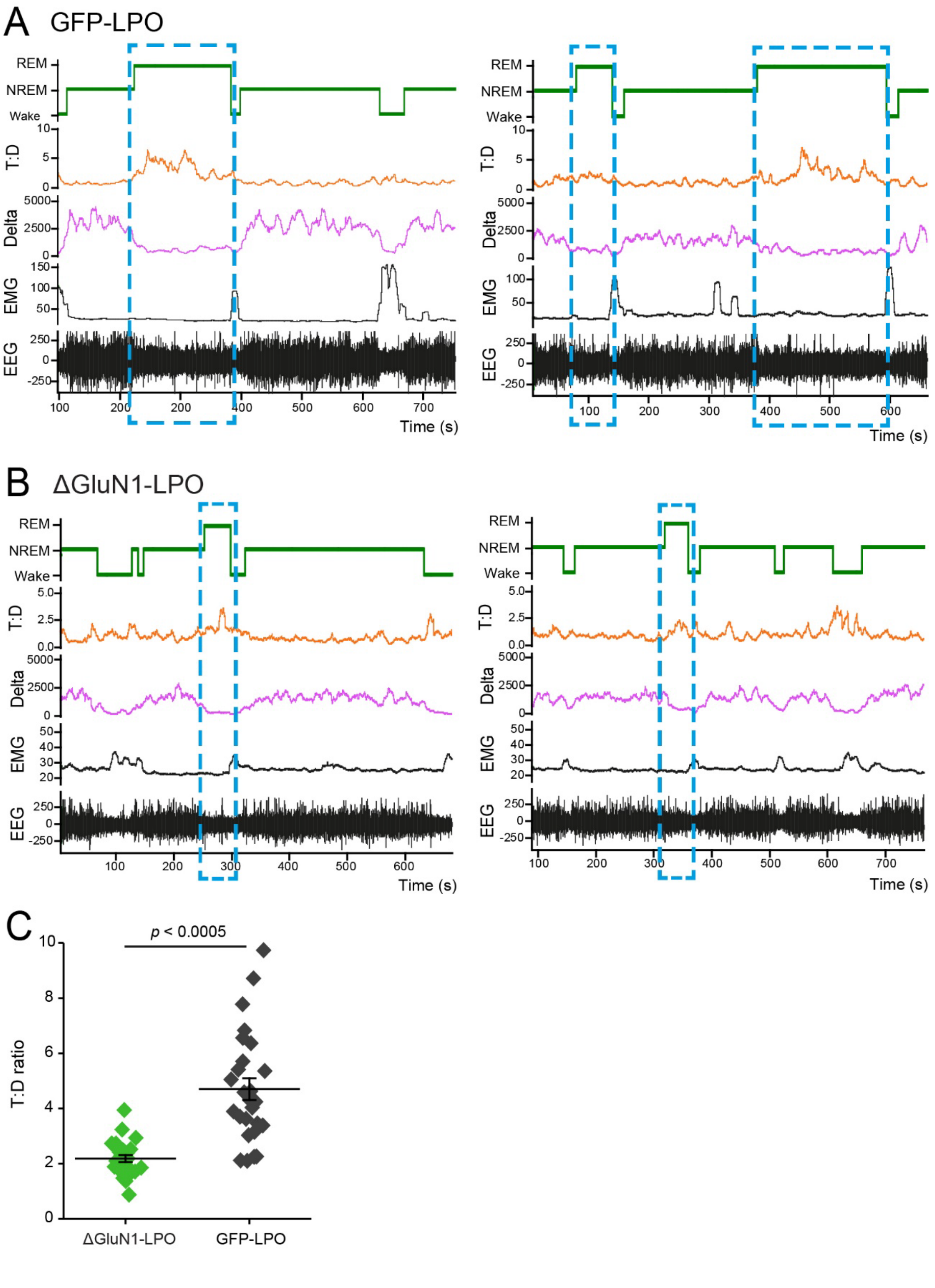
A
GFP-LPO
REM
NREM
Wake
T:D
Delta
EMG
EEG
Time (s)
B
ΔGluN1-LPO
C
p < 0.0005
T:D ratio
ΔGluN1-LPO
GFP-LPO

**Supplementary Figure 3. Deletion of the NR1 subunit from LPO strongly reduces theta waves and the T:D ratio during REM sleep episodes.**

**Supports Figure 3.**

**A** and **B**, examples of sleep recordings from two GFP-LPO (A) and ΔGluN1-LPO (B) animals. From the top: stage, Theta:Delta power ratio (T:D), Delta power, EMG and EEG. Blue dashed rectangles indicate REM sleep episodes. **C**, Quantification of the T:D ratio in ΔGluN1-LPO and GFP-LPO animals. Each point represents a 100s average of the T:D ratio during 3 REM episodes per animal. GFP-LPO, *n*=7; ΔGluN1-LPO, *n*=7, Unpaired Student's *t* test.

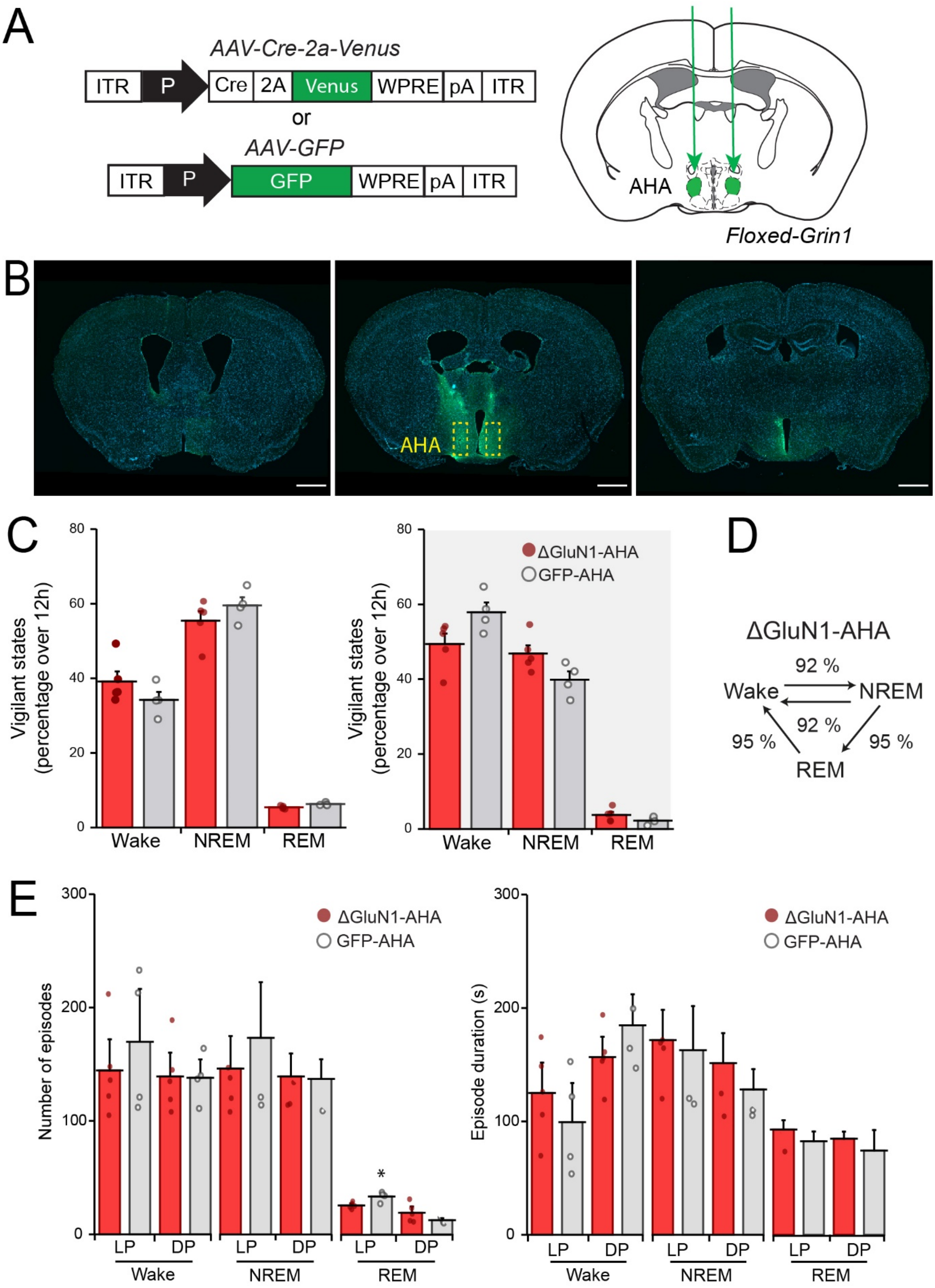
A
AAV-Cre-2a-Venus
ITR P Cre 2A Venus WPRE pA ITR
or
AAV-GFP
ITR P GFP WPRE pA ITR
AHA
Floxed-Grin1
B
AHA
C
Vigilant states (percentage over 12h)
Wake NREM REM
ΔGluN1-AHA
GFP-AHA
D
ΔGluN1-AHA
Wake
NREM
REM
92 %
92 %
95 %
95 %
E
Number of episodes
Episode duration (s)
LP DP
Wake NREM REM

**Supplementary Figure 4. Deletion of the NR1 receptor in a neighbouring area of the hypothalamus to the LPO, the anterior hypothalamic area, does not recapitulate the sleep-wake phenotype of ΔGluN1-LPO mice.**

**Supports Figure 3.**

**A,** *AAV-Cre-2A-Venus* or *AAV-GFP* bilateral injection in the anterior hypothalamic area (AHA), to generate ΔGluN1-AHA and GFP-AHA mice. **B**, viral distribution from LPO to mid-thalamus – the AHA is indicated by the dashed yellow squares. Scale bars, 1 mm. **C**, baseline state distribution shown as % over 12h during LP and DP comparing ΔGluN1-AHA with GFP-AHA mice. 2-way ANOVA and followed by *post-hoc* Sidak's using "virus" and "state" as factors (during LP, "virus": F (1, 21) = 5.393e-006, $p$ = 0.9982; "state": F(2, 21) = 320.9, $p$< 0.0001. During DP, "virus": F (1, 21) = 1.915e-006, $p$ = 0.9989; "state": F(2, 21) = 328.5, $p$< 0.0001). **D**, ΔGluN1-AHA number of transitions between states over 24h represented as % over the control group GFP-AHA. **E**, episode number (left) and mean duration (right) divided by vigilance state and by light and dark period. 2-way ANOVA and followed by *post-hoc* Sidak's using "virus" and "state" as factors. Left panel, during LP "virus": F (1, 21) = 1.453, $p$ = 0.2414; "state": F (2, 21) = 26.89, $p$< 0.0001. During DP "virus": F (1, 21) = 0.1402, $p$ = 0.1402.; "state": F (2, 21) = 85.99, $p$< 0.0001. Right panel during LP, "virus": F (1, 21) = 1.103, $p$ = 0.3056; "state": F (2, 21) = 10.96, $p$< 0.0001. During DP, "virus": F (1, 21) = 0.03044, $p$ = 0.8632; "state": F (2, 21) = 23.95, $p$< 0.0001. GFP-AHA, $n$= 4, ΔGluN1-AHA, $n$= 5. In C and E data are means ± SEM.

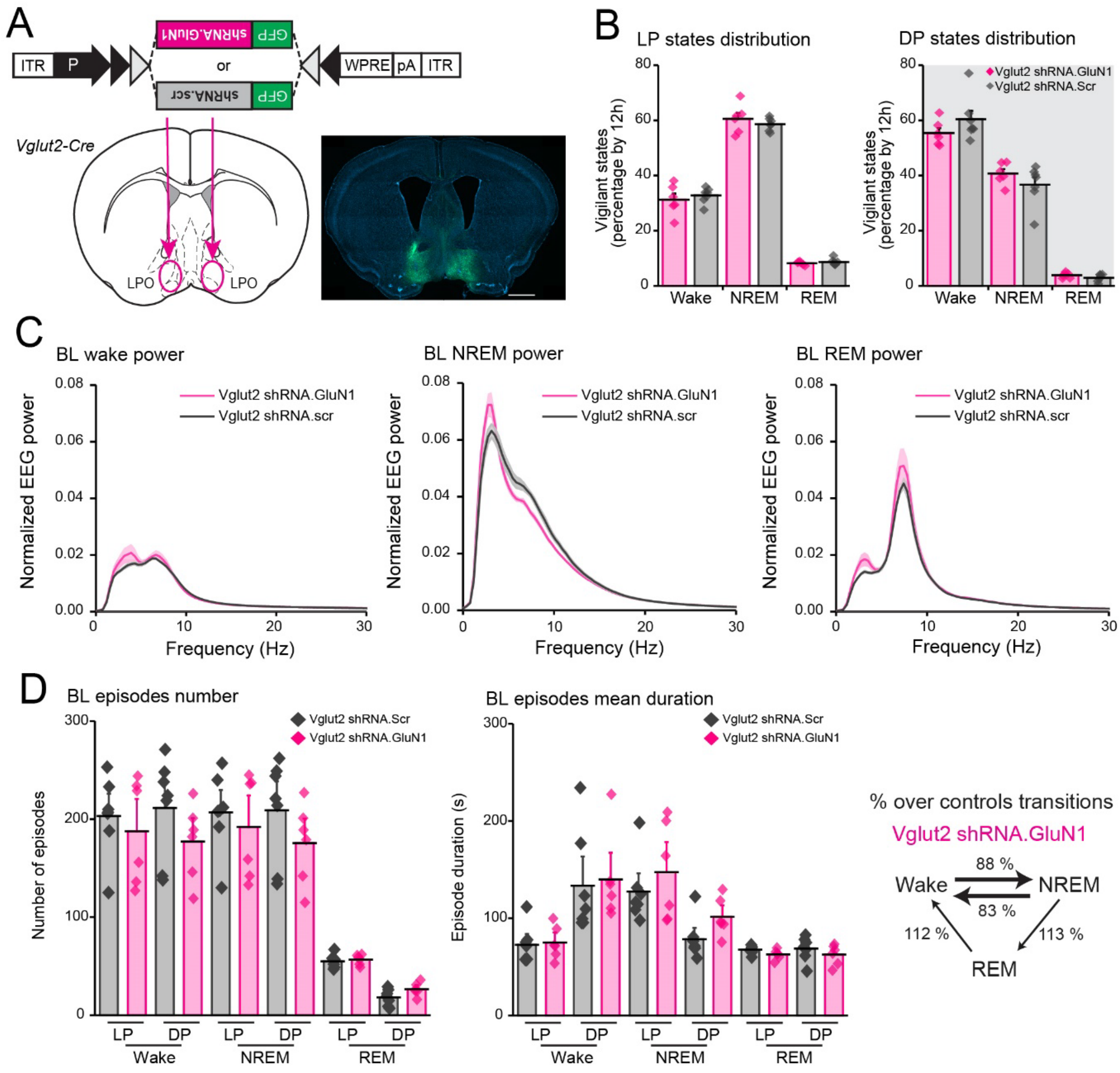
A
ITR
P
shRNA.GluN1
GFP
or
shRNA.scr
WPRE
pA
ITR
Vglut2-Cre
LPO
LPO
B
LP states distribution
DP states distribution
Vigilant states (percentage by 12h)
Vglut2 shRNA.GluN1
Vglut2 shRNA.Scr
Wake
NREM
REM
C
BL wake power
BL NREM power
BL REM power
Normalized EEG power
Vglut2 shRNA.GluN1
Vglut2 shRNA.scr
Frequency (Hz)
D
BL episodes number
Number of episodes
BL episodes mean duration
Episode duration (s)
Vglut2 shRNA.Scr
Vglut2 shRNA.GluN1
LP
DP
Wake
NREM
REM
% over controls transitions
Vglut2 shRNA.GluN1
88 %
83 %
112 %
113 %

**Supplementary Figure 5. Knock-down of the NMDA receptor GluN1 subunit from LPO-Vglut2-expressing neurons does not alter sleep and wake patterns.**

**Supports Figure 6.**

**A**, shRNA-GluN1 or shRNA-scr AAVs were bilaterally injected into the LPO of Vglut2-Cre animals. On the right, immunohistochemistry to map viral vector expression (GFP in green, DAPI in blue). Scale bar, 1mm. **B**, baseline vigilance state amounts calculated as percentage over 12h of the light (left) and dark (right) periods. 2-way ANOVA followed by Sidak's *post-hoc* test using "state" and "virus" as factors. For LP, "state": F (2, 33) = 759.6, P<0.0001; "virus": F (1, 33) = 2.774e-006, P=0.9987. For DP, "state": F (2, 33) = 374.4, P<0.0001; "virus": F (1, 33) = 1.298e-007, P=0.9997. **C**, EEG power spectrum for wake (left), NREM (centre) and REM sleep (right) during 12h light period normalized over the total EEG power. **D** left panel, baseline episode number over 12h comparing Vglut2-shRNA-GluN1 and shRNA-scr animals. 2-way ANOVA followed by Sidak's *post-hoc* test using "state" and "virus" as factors ("state": F (5, 66) = 58.35, P<0.0001; "virus": F (1, 66) = 2.844, P=0.0964); centre panel, episodes mean duration by 12h during baseline recordings. 2 way ANOVA followed by Sidak's *post-hoc* test using "state" and "virus" as factors ("state": F (5, 66) = 17.58, P<0.0001; "virus": F (1, 66) = 1.044, P=0.3106); right panel, Vglut2-shRNA-GluN1 number of transitions between states represented as percentage over the Vglut2-shRNA-scr baseline. Vglut2-shRNA-GluN1, *n* = 6; Vglut2-shRNA-scr, *n* = 7. In B, C and D data are represented as mean ± SEM.

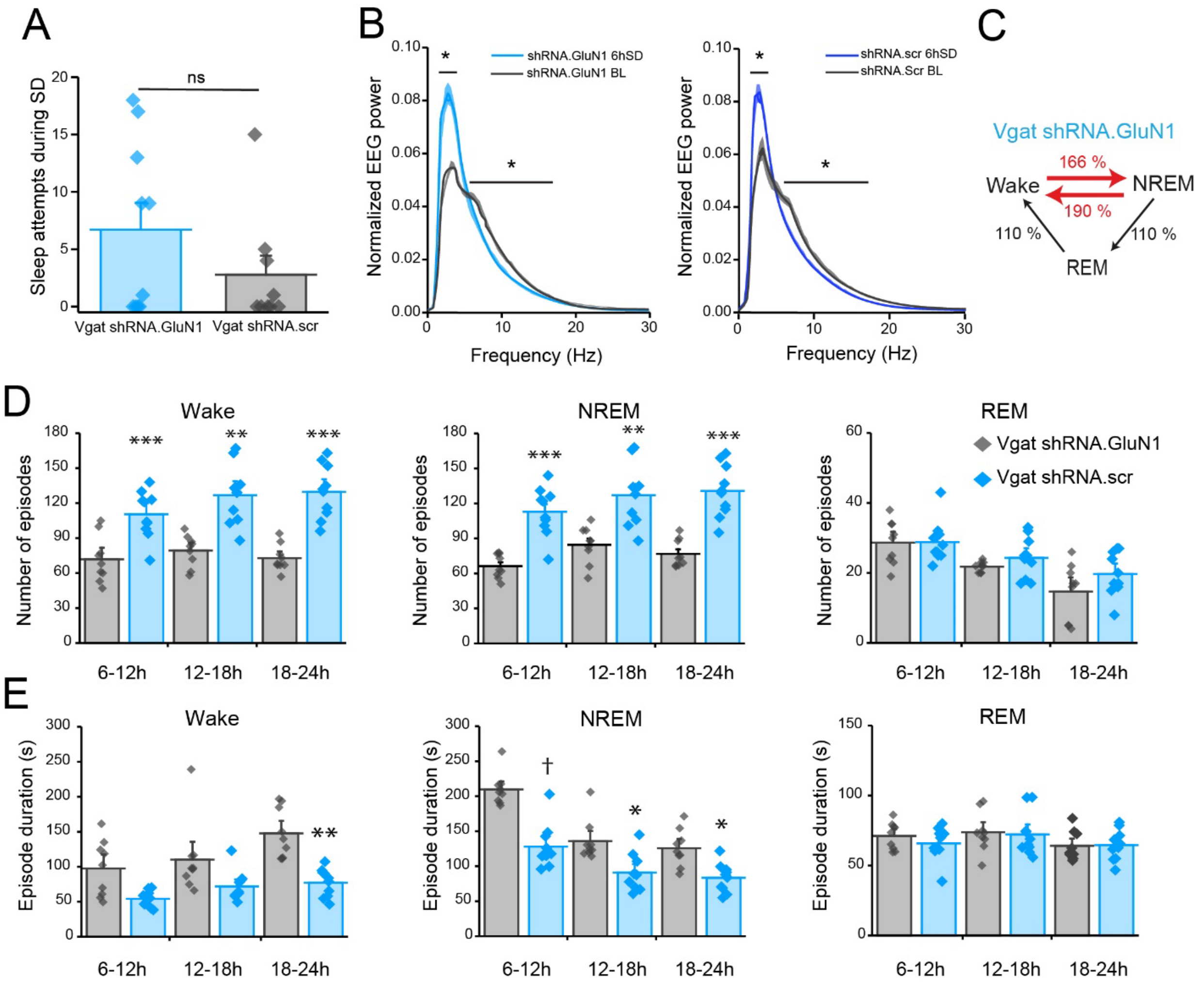

A
Sleep attempts during SD
ns
Vgat shRNA.GluN1
Vgat shRNA.scr
B
Normalized EEG power
shRNA.GluN1 6hSD
shRNA.GluN1 BL
shRNA.scr 6hSD
shRNA.Scr BL
Frequency (Hz)
C
Vgat shRNA.GluN1
166 %
190 %
Wake
NREM
REM
110 %
110 %
D
Wake
NREM
REM
Number of episodes
Vgat shRNA.GluN1
Vgat shRNA.scr
6-12h
12-18h
18-24h
E
Episode duration (s)

**Supplementary Figure 6. Vgat-shRNA-GluN1 animals maintain sleep fragmentation even after 6h sleep deprivation.**

**Supports Figure 6.**

**A**, number of NREM sleep attempts during 6h sleep deprivation comparing Vgat- shRNA-GluN1 animals to Vgat-shRNA-scr ones. Kolmogorov-Smirnov test (P = 0.4708). **B**, NREM EEG power spectrum comparing 1h after sleep deprivation to the same circadian time during baseline recordings, to represent delta power rebound in Vgat-shRNA-GluN1 (left) and Vgat-shRNA-scr (right) animals. 2-way ANOVA followed by Sidak's *post-hoc* test using "virus" and "frequency" as factors. For Vgat-shRNA-GluN1, "frequency": F (89, 1620) = 613.6, P<0.0001; "virus": F (1, 1620) = 1.075, P=0.2999. For Vgat-shRNA-scr, "frequency": F (89, 1620) = 623.7, P<0.0001; "virus": F (1, 1620) = 2.081, P=0.1493. **C**, Vgat-shRNA-GluN1 animals number of transitions during the 18h following SD represented as a percentage over the number of transitions exhibited by Vgat-shRNA-scr control animals. **D**, episode number during wake (left), NREM (centre) and REM sleep (right) following 6hSD. 2-way RM ANOVA followed by Sidak's *post-hoc* test using "state" and "virus" as factors ("state": F (2.537, 43.13) = 217.9, P<0.0001; "virus": F (1, 17) = 37.43, P<0.0001). **E**, episode mean duration following 6h SD for wake (left), NREM (centre) and REM sleep (right). 2-way RM ANOVA followed by Sidak's *post-hoc* test using "state" and "virus" as factors ("state": F (4.785, 81.35) = 55.57, P<0.0001; "virus": F (1, 17) = 40.01, P<0.0001). Vgat-shRNA-GluN1, *n* = 10; Vgat-shRNA-scr, *n* = 9. Data in A, B, D and E are means ± SEM. In B, D and E, *P* < 0.05, ** *P*< 0.005, *** *P*< 0.0005, † *P*< 0.00005 from multiple comparisons analysis corrected by Sidak's *post-hoc* test.
